# Palmitoylation regulates neuropilin-2 localization and function in cortical neurons and conveys specificity to semaphorin signaling via palmitoyl acyltransferases

**DOI:** 10.1101/2022.10.28.514228

**Authors:** Eleftheria Koropouli, Qiang Wang, Rebeca Mejias, Randal Hand, Tao Wang, David D. Ginty, Alex L. Kolodkin

## Abstract

Secreted semaphorin 3F (Sema3F) and semaphorin 3A (Sema3A) exhibit remarkably distinct effects on deep layer excitatory cortical pyramidal neurons; Sema3F mediates dendritic spine pruning, whereas Sema3A promotes the elaboration of basal dendrites. Sema3F and Sema3A signal through distinct holoreceptors that include neuropilin-2 (Nrp-2)/plexinA3 (PlexA3) and neuropilin-1 (Nrp-1)/PlexA4, respectively. We find that Nrp-2 and Nrp-1 are S-palmitoylated in cortical neurons and that palmitoylation of select Nrp-2 cysteines is required for its proper subcellular localization and also for Sema3F/Nrp-2-dependent dendritic spine pruning in cortical neurons, both in vitro and in vivo. Moreover, we show that the palmitoyl acyltransferase DHHC15 is required for Nrp-2 palmitoylation and Sema3F/Nrp-2-dependent dendritic spine pruning, but it is dispensable for Nrp-1 palmitoylation and Sema3A/Nrp-1-dependent basal dendritic elaboration. Therefore, palmitoyl acyltransferase-substrate specificity is essential for establishing compartmentalized neuronal structure and functional responses to extrinsic guidance cues.

**HIGHLIGHTS:**

- Neuropilins (Nrps) are S-palmitoylated in vitro and in vivo in the central nervous system
- S-palmitoylation of select Nrp-2 cysteines confers subcellular localization specificity and is required for semaphorin 3F-dependent dendritic spine pruning in cortical neurons
- Distinct palmitoyl acyltransferases mediate Nrp-2 and Nrp-1 palmitoylation and function, imparting specificity to semaphorin signaling

## INTRODUCTION

The central nervous system (CNS) consists of numerous disparate classes of neurons with remarkably distinct morphologies and functions. This is particularly prominent in laminated structures, including the cerebral cortex, where pyramidal neurons occupying different cortical layers acquire distinct morphologies and their processes exhibit subcellular compartmentalization that mediates distinct functions (Spruston, 2008). For example, dendritic spines, which receive the vast majority of excitatory inputs, have specific subcellular distributions that directly impact electrical properties of neurons and subsequently the activity within neuronal circuits. Numerous studies highlight the physiological importance of dendritic spines, linking alterations in spine number and morphology to various neuropsychiatric disorders (Penzes et al., 2011).

One class of proteins implicated in nervous system development are semaphorins (Koropouli and Kolodkin, 2014). Class 3 secreted semaphorins semaphorin 3F (Sema3F) and semaphorin 3A (Sema3A) play critical roles in several aspects of neural development and function, including axon guidance, axon pruning, dendritic arborization, dendritic spine distribution, synaptic transmission and homeostatic synaptic plasticity (Danelon et al., 2020; Demyanenko et al., 2014; Gu et al., 2003; Koropouli and Kolodkin, 2014; Li et al., 2022; Riccomagno et al., 2012; Sahay et al., 2005; Tran et al., 2009; Wang et al., 2017). In mammalian cortical neurons, Sema3F mediates pruning of excess dendritic spines along the apical dendrite of layer V cortical pyramidal neurons, whereas Sema3A promotes the elaboration of basal dendrites in the same neurons (Tran et al., 2009). Sema3F and Sema3A bind distinct holoreceptor complexes that include neuropilin (Nrp) and plexin (Plex) transmembrane proteins; Sema3F exerts many of its effects via a holoreceptor complex that includes Nrp-2/PlexA3, whereas Sema3A acts through a holoreceptor complex that includes Nrp-1/PlexA4 (Yaron et al., 2005). Recent work provides insight into the signaling pathways that mediate Sema3F/Nrp-2-dependent cytoskeletal rearrangements resulting in dendritic spine pruning in cortical neurons, including contributions by specific immunoglobin superfamily transmembrane proteins that mediate select responses to secreted semaphorins and proteins that regulate actin cytoskeleton dynamics (Demyanenko et al., 2014; Duncan et al., 2021). However, molecular mechanisms that regulate neuropilin subcellular localization and underlie divergent Sema3F and Sema3A functions in cortical neurons remain largely unknown.

The attachment of the fatty acid palmitate on thiol groups of cysteine residues, known as S-palmitoylation (hereafter referred to as palmitoylation), is a reversible posttranslational modification that dynamically regulates protein localization and function of a vast protein repertoire in different tissues (Linder and Deschenes, 2007; Salaun et al., 2010). In the nervous system, palmitoylation is critically involved in all aspects of neural development and function (Fukata and Fukata, 2010; Kang et al., 2008), including axon outgrowth (Tortosa et al., 2017), dendritic arborization (Takemoto-Kimura et al., 2007), spine formation (George et al., 2015; Kang et al., 2008; Kutzleb et al., 1998), synapse assembly, synaptic transmission (El-Husseini et al., 2002; Hayashi et al., 2005; Keith et al., 2012; Lin et al., 2009; Sanders et al., 2020; Thomas et al., 2012) and synaptic plasticity (Brigidi et al., 2014). Strikingly, there is an apparent interaction between palmitoylation and synaptic transmission since neuronal activity can regulate protein palmitoylation (Brigidi et al., 2014; Hayashi et al., 2005; Kang et al., 2008). Palmitoylation is catalyzed by palmitoyl acyltransferases, enzymes that harbor the catalytically active Asp-His-His-Cys (DHHC) signature motif (thereby, also referred to as DHHCs). Originally discovered in the yeast *Saccharomyces cerevisiae* (Lobo et al., 2002; Roth et al., 2002), thus far 23 DHHC enzymes are predicted to exist in humans and mice. DHHCs exhibit overlapping, yet distinct, specificities for their substrates (Huang et al., 2004; Roth et al., 2006). Despite systematic and extensive efforts (Roth et al., 2006), our knowledge of DHHCs substrates remains very limited, hindering our understanding of the roles that DHHCs play in neuronal development and function.

Here, we show that both Nrp-2 and Nrp-1 are palmitoylated in cortical neurons in vitro and in vivo, and that palmitoylation of select Nrp-2 cysteine residues is required for correct Nrp-2 subcellular localization and for Sema3F/Nrp-2-dependent dendritic spine pruning both in vitro and in vivo. Our findings also reveal that the palmitoyl acyltransferase DHHC15 is required for proper Nrp-2 palmitoylation and Sema3F/Nrp-2-dependent dendritic spine pruning in layer V cortical pyramidal neurons, but not for Nrp-1 palmitoylation or for Sema3A/Nrp-1-dependent basal dendrite elaboration in these same neurons. These results highlight the importance of guidance cue receptor posttranslational lipid modifications to regulate distinct aspects of cortical neuron morphology.

## RESULTS

### Nrp-2 and Nrp-1 exhibit distinct cell surface distribution patterns and global inhibition of palmitoylation leads to Nrp-2 mislocalization

To understand the mechanisms by which Sema3F and Sema3A exert distinct effects on the development of layer V cortical pyramidal neurons, we investigated the localization and function of their obligate co-receptors, Nrp-2 and Nrp-1, respectively. First, we assessed the localization of Nrp-2 and Nrp-1 on the surface of COS-7 cells owing to the well-articulated plasma membrane and large circumference of these cells in culture. COS-7 cells transfected with flag-tagged wild-type Nrp-2 or Nrp-1 expression plasmids were subjected to surface staining with a flag antibody to visualize Nrp localization on the plasma membrane. We observed that Nrp-2 is distributed on the COS-7 cell surface in a clustered pattern consisting of numerous discrete puncta (Figure 1A), whereas Nrp-1 is evenly distributed over the entire plasma membrane with little evidence of clustering (Figure 1B). Particle analysis (see Materials and Methods section) provides a quantitative assessment of protein clustering and reveals a marked difference between the membrane localization of Nrp-2 and Nrp-1 (Figure 1C). This robust Nrp-2 clustering is reminiscent of the cell surface clustering displayed by palmitoylated proteins in COS-7 cells (Webb et al., 2000).

**Figure 1.**
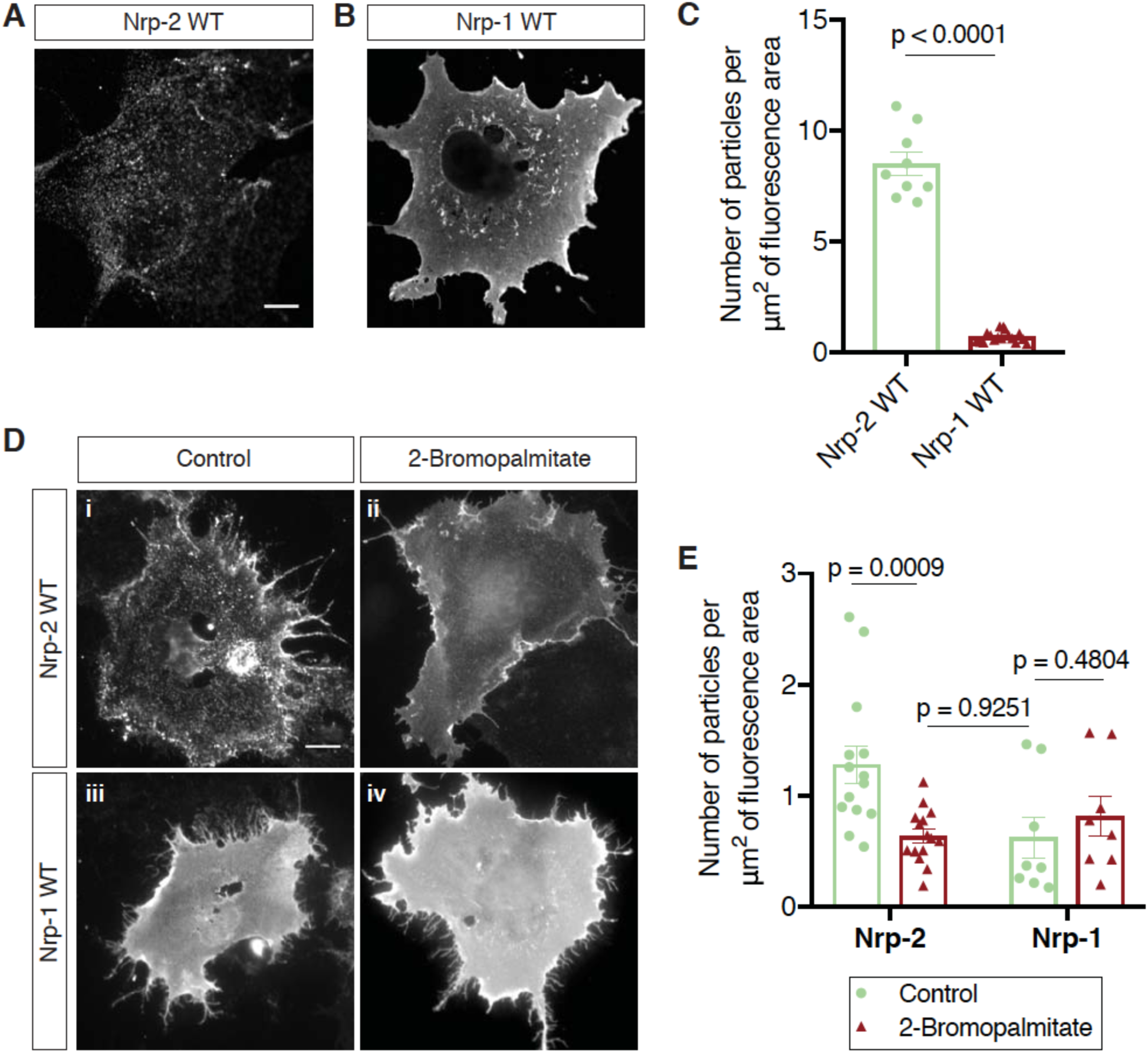
Distinct cell surface localization patterns of Nrp-2 and Nrp-1 are abolished by global inhibition of palmitoylation. **(A-C)** Surface localization of Nrp-2 and Nrp-1 in COS-7 cells expressing exogenous flag-tagged Nrp-2 wild-type (WT) (A) or Nrp-1 WT (B). Cells were subjected to surface staining with a flag antibody to visualize cell surface protein. Representative images are shown for each protein. Nrp-2 appears clustered, whereas Nrp-1 appears diffusely localized on the cell surface. Scale bar, 12 µm. (C) Quantification of protein clustering, measured as the number of particles per µm^2^ of fluorescence area (Clustering analysis, see Materials and Methods). Data are plotted in a scatter dot plot with mean ± SEM (SEM, Standard Error of the Mean). Two-tailed t test, Nrp-2, n = 9; Nrp-1, n = 17. **(D, E)** Effects of 2-bromopalmitate on cell surface Nrp localization in heterologous cells. COS-7 cells expressing exogenous flag-tagged WT Nrp-2 or Nrp-1 were treated overnight with medium containing either 10 µM 2-bromopalmitate (ii, iv) or the same concentration of solvent (i, iii). Cells were subjected to surface staining with a flag antibody. Representative images are shown for each protein. Upon control treatment, Nrp-2 appears highly clustered (i), while Nrp-1 has an even diffuse distribution on the plasma membrane (iii). Upon treatment with 2-bromopalmitate, Nrp-2 assumes diffuse distribution (ii), similar to Nrp-1 (iii, iv). Scale bar, 20 µm. (E) Quantification of protein clustering shown in (D), as mentioned above. Data are plotted in a scatter dot plot with mean ± SEM. Two-tailed t test; Nrp-2 control, n = 14; Nrp-2 2-bromopalmitate, n = 15; Nrp-1 control, n = 8; Nrp-1 2-bromopalmitate, n = 8, where n is the number of cells analyzed. The online version of this article includes the following source data for figure 1: **Source data 1.** Raw data for Figure 1C. **Source data 2.** Raw data for Figure 1E.

To explore the potential role of palmitoylation in Nrp-2 surface localization we used a pharmacological approach employing 2-bromopalmitate, which is a specific and irreversible inhibitor of protein palmitoylation (Jennings et al., 2009; Resh, 2006). We treated COS-7 cells expressing flag-tagged wild-type Nrp-2 (Nrp-2^wild-type^) with 2-bromopalmitate, or a control solution, and visualized cell surface Nrp-2. Following bath application of 2-bromopalmitate, Nrp-2 was markedly redistributed compared to the controls, displaying a diffuse localization on the cell surface similar to Nrp-1, which is localized diffusely following either control or 2-bromopalmitate treatment (Figures 1D and 1E). 2-bromopalmitate-induced Nrp-2 dispersion is consistent with previous reports showing 2-bromopalmitate-induced diffusion of otherwise clustered palmitoylated proteins in non-neuronal cell lines (Webb et al., 2000) and in neurons (El-Husseini et al., 2002). These results show that Nrp-2 and Nrp-1 exhibit distinct cell surface compartmental localization and that protein palmitoylation is required for Nrp-2 cell surface clustering in heterologous cells in vitro.

### Nrp-2 and Nrp-1 are palmitoylated in cortical neurons in vitro and in the mouse brain, exhibiting overlapping palmitoylation patterns

Given the significant Nrp-2 cell surface distribution perturbations we observed upon 2-bromopalmitate bath application to COS-7 cells, we next examined whether Nrps are palmitoylated. Both Nrp-2 and Nrp-1 harbor cysteine residues in their transmembrane and membrane-proximal (juxtamembrane) domains (Figure 2A, in red). Nrp-2 also has a lone cysteine residue, C897 (Figure 2A, in red), carboxy-terminal (C-terminal) to the transmembrane domain and two adjacent C-terminal cysteine residues (Figure 2A, in green). These Nrp-2 and Nrp-1 cysteine residues are all highly phylogenetically conserved among vertebrate species (Figure 2—figure supplement 1). The location and conservation of these cysteine residues, coupled with the presence of palmitoyl acceptor amino acid residues predicted using CSS-Palm software (Ren et al., 2008), suggested that these cysteine residues are palmitoyl acceptor sites. To explore this further, we used the Acyl-Biotin Exchange (ABE) assay, a well-established biochemical approach for the detection of palmitoylation on thiol groups of cysteines (S-palmitoylation) (Drisdel and Green, 2004; Kang et al., 2008; Roth et al., 2006; Wan et al., 2007). In this assay, we used the postsynaptic density protein 95 (PSD-95) as a positive palmitoylation control because it is palmitoylated (Topinka and Bredt, 1998), and synapse-associated protein 102 (SAP102) as a negative palmitoylation control because it is not (Kang et al., 2008). We observed that both Nrp-2 and Nrp-1 isolated from adult mouse forebrain and from embryonic day 14.5 (E14.5) mouse cortex are palmitoylated (Figure 2B), as are both Nrps isolated from cortical neurons 28 days in vitro (DIV) derived from E14.5 embryos (Figure 2C). The detection of Nrp palmitoylation from early embryonic stages (E14.5) to adulthood raises the possibility that palmitoylation of these co-receptors is required throughout neural development and in the adult to regulate Nrp trafficking and function.

**Figure 2.**
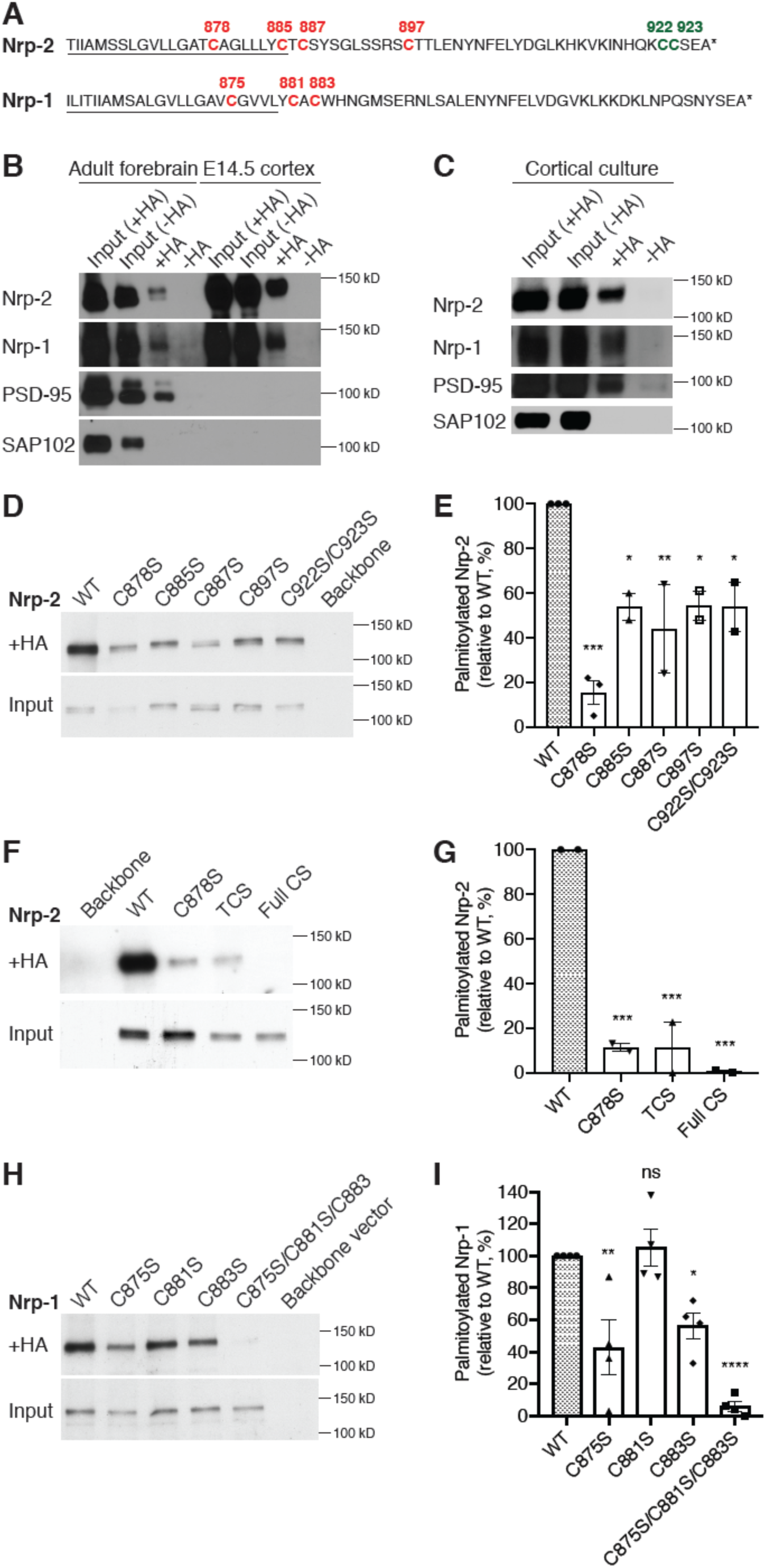
Neuropilins are palmitoylated in deep layer primary cortical neurons and in the mouse brain. **(A)** Amino acid sequence of transmembrane (underlined) and cytoplasmic domains of Nrp-2 and Nrp-1, as predicted by Ensembl genome database. Both Nrp-2 and Nrp-1 harbor cysteine residues in their transmembrane and juxtamembrane domains (depicted as red), whereas Nrp-2 also possesses a C-terminal di-cysteine motif (depicted as green). Cysteine numeration accords with their position in the amino acid sequence of *Mus musculus* Nrp2 (Nrp2-202, isoform A17, ENSMUST00000063594.13) for Nrp-2 and *Mus musculus* Nrp-1 (Nrp-1-201, ENSMUST00000026917.10) for Nrp-1. **(B, C)** Acyl-Biotin Exchange (ABE) performed on adult mouse forebrain and E14.5 cortex (B) or on E14.5 DIV28 cortical neuron cultures (C). Hydroxylamine-treated (+HA) samples show palmitoylated protein, while -HA samples serve as a negative control. Both Nrp-2 and Nrp-1 are palmitoylated in the mouse brain and cortex, in vivo and in vitro, similar to PSD-95 that serves as a positive palmitoylation control. SAP102, known not to be palmitoylated, serves as a negative control. **(D, E)** ABE on Neuro2A cells expressing flag-tagged Nrp-2 wild-type (WT) or CS point mutants. (D) Nrp-2 immunoblots show palmitoylated (+HA) and input samples. –HA samples (not shown) were included in the experiment. (E) Quantification of palmitoylated protein levels (fraction of the protein that is palmitoylated) calculated as the ratio of +HA to the respective input and plotted in a scatter dot plot with mean ± SEM (n = 2-3 experiments). Palmitolyated WT Nrp-2 is set at 100% and CS mutants are expressed as a percentage of WT. The cysteine residue C878 stands out as a major Nrp-2 palmitate acceptor site. One-way ANOVA followed by Dunnett’s test for multiple comparisons (Nrp-2 CS mutants are compared to Nrp-2 WT (set at 100%): C878S, p = 0.0002; C885S, p = 0.0189; C887S, p = 0.0062; C897S, p = 0.0200; C922S/C923S, p = 0.0189). **(F, G)** ABE on Neuro2A cells expressing flag-tagged Nrp-2 wild-type (WT) or various CS mutants. (F) Nrp-2 immunoblots show palmitoylated (+HA) and input samples. –HA samples (not shown) were included in the experiment. (G) Quantification of palmitoylated protein levels (as explained above), plotted in scatter dot plots with mean ± SEM (n = 2 experiments). The Nrp-2 transmembrane and membrane-proximal cysteines are major palmitoylation sites. One-way ANOVA followed by Dunnett’s test for multiple comparisons (Nrp-2 CS mutants are compared to Nrp-2 WT (set at 100%): C878S, p = 0.0009; TCS, p = 0.0009; Full CS, p = 0.0006). **(H, I)** ABE on Neuro2A cells expressing flag-tagged Nrp-1 wild-type (WT) or CS point mutants. (H) Nrp-1 immunoblots show palmitoylated (+HA) and input samples. (I) Quantification of palmitoylated protein levels (as explained above), plotted in scatter dot plot including mean ± SEM (n = 4 experiments). Nrp-1 is palmitoylated mostly on cysteines C875 and C883. One-way ANOVA followed by Dunnett’s test for multiple comparisons (Nrp-1 CS mutants are compared to Nrp-1 WT (set at 100%): C875S, p = 0.0042; C881S, p = 0.9877; C883S, p = 0.0272; C875S/C881S/C883S, p < 0.0001; ns, not significant). The online version of this article includes the following source data and figure supplement(s) for figure 2: **Source data 1.** Raw, unedited blot from Figure 2B. **Source data 2.** Raw, unedited blot from Figure 2B. **Source data 3.** Raw, unedited blot from Figure 2B. **Source data 4.** Raw, labeled blot from Figure 2B. **Source data 5.** Raw, labeled blot from Figure 2B. **Source data 6.** Raw, labeled blot from Figure 2B. **Source data 7.** Raw, unedited blot from Figure 2C. **Source data 8.** Raw, unedited blot from Figure 2C. **Source data 9.** Raw, unedited blot from Figure 2C. **Source data 10.** Raw, unedited blot from Figure 2C. **Source data 11.** Raw, labeled blot from Figure 2C. **Source data 12.** Raw, labeled blot from Figure 2C. **Source data 13.** Raw, labeled blot from Figure 2C. **Source data 14.** Raw, labeled blot from Figure 2C. **Source data 15.** Raw, unedited blot from Figure 2D. **Source data 16.** Raw, labeled blot from Figure 2D. **Source data 17.** Raw data for Figure 2E. **Source data 18.** Raw, unedited blot from Figure 2F. **Source data 19.** Raw, unedited blot from Figure 2F. **Source data 20.** Raw, labeled blot from Figure 2F. **Source data 21.** Raw, labeled blot from Figure 2F. **Source data 22.** Raw data for Figure 2G. **Source data 23.** Raw, unedited blot from Figure 2H. **Source data 24.** Raw, labeled blot from Figure 2H. **Source data 25.** Raw data for Figure 2I. **Figure supplement 1.** Conserved cysteine residues lie in the transmembrane and cytoplasmic domains of neuropilins.

We next sought to identify the Nrp-2 and Nrp-1 cysteine residues that serve as palmitate acceptors. For this analysis, we generated point mutations that replaced individual Nrp-2 and Nrp-1 cysteines with a serine residue, which cannot be S-palmitoylated. Cysteine-to-serine (CS) point mutant Nrp expression plasmids were transfected into neuroblastoma-2a (Neuro2A) cells on which the ABE assay was performed. This analysis revealed that Nrp-2 is palmitoylated, albeit to variable extents, on all transmembrane and membrane-proximal cysteines as well as on the C-terminal di-cysteine motif (Figures 2D and 2E). The triple CS (TCS) Nrp-2 mutant C878S/C885S/C887S (*Nrp-2^TCS^*: substituting serine for the three transmembrane/juxtamembrane cysteines) and the Full CS Nrp-2 mutant C878S/C885S/C887S/C897S/C922S/C923S (*Nrp-2^Full^ ^CS^*: substituting all transmembrane and cytoplasmic cysteines) display very little to no palmitoylation, respectively (Figures 2F and 2G). A similar analysis for Nrp-1 revealed that Nrp-1 is palmitoylated on cysteines C875 and C883, but apparently not on C881 (Figures 2H and 2I). These results show that Nrp-2 and Nrp-1 display similar palmitoylation patterns in their transmembrane and membrane-proximal segments, and that Nrp-2 also harbors C-terminal palmitoylation sites that are not present in Nrp-1.

### Select palmitoyl acceptor cysteines regulate the localization and trafficking of Nrp-2 across subcellular compartments

Palmitoylation enables proteins to be anchored onto specialized membrane domains, to interact with other proteins, and to shuttle between the plasma membrane and intracellular organelles (Salaun et al., 2010). The compartmentalized distribution of Nrp-2 on the cell surface of COS-7 cells in culture, which is palmitoylation-dependent (Figures 1D and 1E), prompted us to investigate the role of palmitoylation in Nrp-2 localization, trafficking, and function in cortical neurons.

First, we addressed the role of palmitoyl acceptor Nrp-2 cysteines in Nrp-2 surface localization in cell lines. We expressed wild-type or various CS mutant flag-tagged Nrp-2 proteins in COS-7 cells and assessed their distribution with surface staining. This assay revealed that localization of Nrp-2^C922S/C923S^, which lacks the C-terminal di-cysteine motif, is for the most part punctate with little difference from Nrp-2^WT^, whereas Nrp-2^TCS^ and Nrp-2^Full^ ^CS^ proteins are profoundly mislocalized (Figure 3—figure supplement 1A-1C), exhibiting a non-patterned distribution over the entire plasma membrane that resembles Nrp-1. For our localization analysis experiments we also generated pHluorin-tagged Nrp-2 mutants. The performance of the same localization assay with pHluorin-tagged Nrp-2 wild-type and CS mutants yielded similar results (Figures 3A and 3B). The diffuse distribution of palmitoylation-deficient Nrp-2 CS mutants on the cell surface is reminiscent of Nrp-2^WT^ protein distribution following treatment with 2-bromopalmitate (Figure 1D). Further, these results are in line with the palmitoylation patterns we observed for these same Nrp-2 CS mutants (Figures 2D-2G).

**Figure 3.**
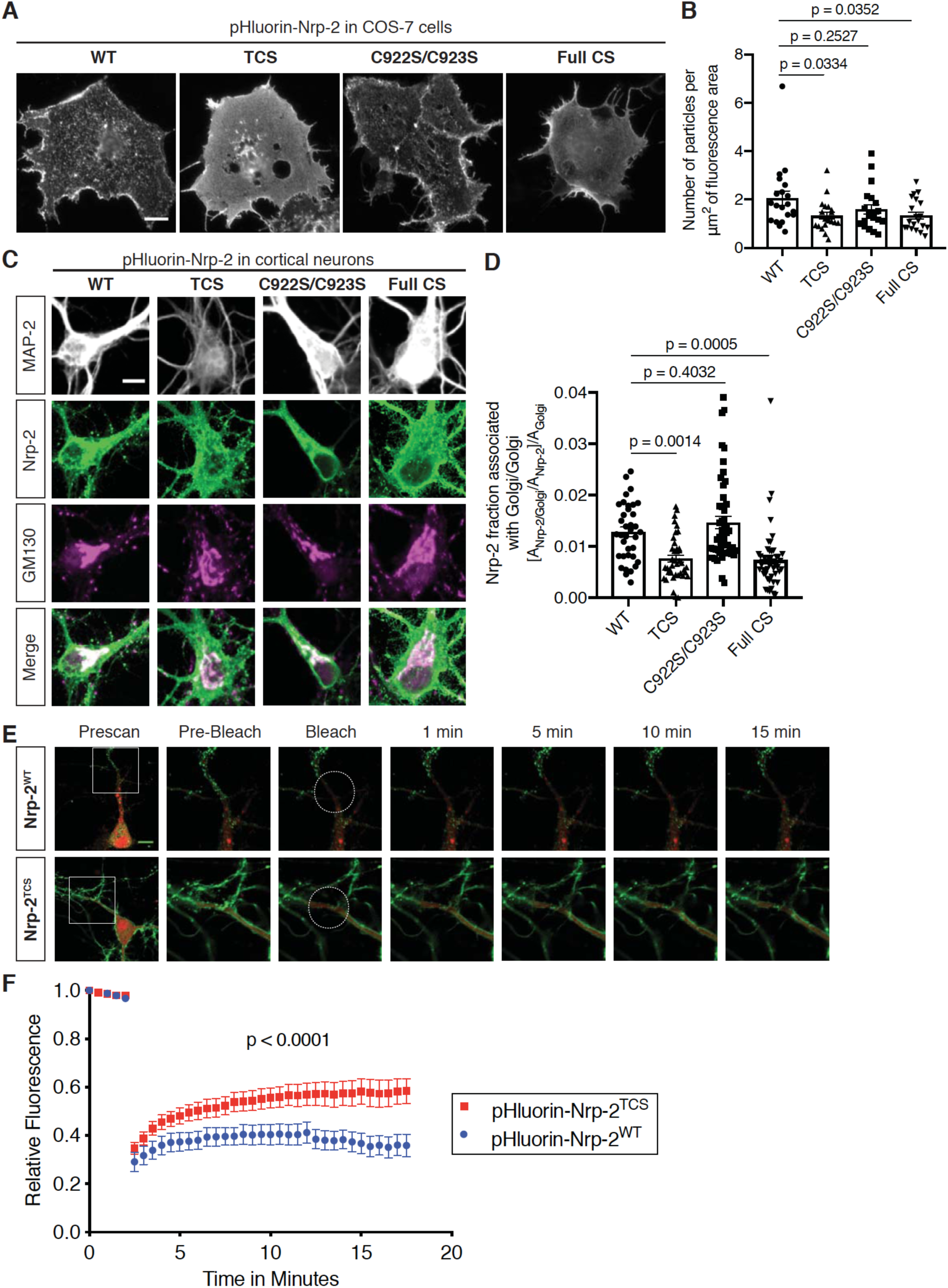
Differential requirements for distinct Nrp-2 palmitoyl acceptor cysteines in regulating subcellular Nrp-2 localization and trafficking. **(A, B)** Effects of Nrp-2 cysteines on Nrp-2 protein cell surface localization in heterologous cells. (A) Panels show COS-7 cells expressing pHluorin-tagged Nrp-2 wild-type (WT) or various CS mutants and subjected to Nrp-2 surface staining with a GFP antibody. Scale bar, 15 µm. (B) Quantification of protein clustering with particle analysis (as mentioned earlier), presented as number of particles per µm^2^ of fluorescence area and plotted in scatter dot plot including mean ± SEM. WT and C922S/C923S Nrp-2 proteins are distributed in the form of smaller particles (puncta), whereas TCS and Full CS Nrp-2 proteins localize on the surface as large protein clusters. One-way ANOVA followed by Dunnett’s test for multiple comparisons; n = 20 cells for each plasmid. (**C, D**) Effects of Nrp-2 cysteines on Nrp-2 localization at the Golgi apparatus in cortical neuron cultures. Colocalization of pHluorin-tagged Nrp-2 WT or cysteine mutants with the Golgi apparatus marker GM130, in E14.5 DIV17 *Nrp-2^-/-^* primary cortical neurons. (**C**) Representative images of neurons expressing different Nrp-2 proteins (WT and CS point mutants). In merged panels, the regions where Nrp-2 and GM130 puncta of similar intensity colocalize appear white. Nrp-2 WT and C922S/C923S exhibit very strong association with Golgi cisternae. By contrast, Nrp-2 TCS or Full CS colocalize with the Golgi to a significantly lesser extent. Of note, following Nrp-2 staining, neurons expressing Nrp-2^Full^ ^CS^ appear “hairy” (right-most column, second row); this is a common phenotype of this Nrp-2 CS mutant, indicative of its diffuse distribution in all membranes including filopodia. Scale bar, 7 µm. (**D**) Quantification of the degree of colocalization between Nrp-2 and GM130, plotted as the fraction of Nrp-2 associated with Golgi [A_Nrp-2/Golgi_/A_Nrp-2_] normalized to the quantity of Golgi (Golgi Area, A_Golgi_) present in each neuron ([A_Nrp-2/Golgi_/A_Nrp-2_]/A_Golgi_) (see Materials and Methods). Columns show pooled data plotted in scatter dot plot including mean ± SEM. One-way ANOVA (p < 0.0001) followed by Dunnett’s test for multiple comparisons. WT, n = 36; TCS, n = 40; C922S/C923S, n = 50; Full CS, n = 50, where n is the number of neurons analyzed. (**E, F**) Fluorescence Recovery After Photobleaching (FRAP) analysis on E14.5 DIV10 wild-type cortical neurons expressing pHluorin-tagged *Nrp-2-ires-dsRED* plasmids. (**E**) Time-lapse image sequences are shown for each Nrp-2 protein. Pre-bleach and post-bleach panels depict the areas surrounded by white squares in prescan panels. White dashed circles delineate the region of interest (ROI) selected for photobleaching. Note the higher diffusibility of Nrp-2^TCS^ compared to Nrp-2^WT^ and the difference in surface distribution of WT and TCS Nrp-2, which appear clustered and diffuse respectively. Scale bar, 10 µm. (F) Quantitative analysis of fluorescence recovery kinetics after photobleaching. Pooled data are plotted as mean ± SEM. Extra sum-of-squares F test (WT, n = 7 neurons, Rmax=0.4463, T1/2=1.242e-016; TCS, n = 8 neurons, Rmax=0.5824, T1/2=1.348e-016, where Rmax is maximum recovery). The online version of this article includes the following source data and figure supplement(s) for figure 3: Source data 1. Raw data for Figure 3B. Source data 2. Raw data for Figure 3D. Source data 3. Raw data for Figure 3F. Figure supplement 1. Distinct requirements for Nrp-2 palmitoyl acceptor cysteines in Nrp-2 cell surface distribution in COS-7 cells. Figure supplement 1—source data 1. Raw data for Figure 3*—figure supplement 1B and 1C*. Figure supplement 2. Severe defects in the cell surface localization of palmitoylation-deficient Nrp-2 in primary cortical neurons. Figure supplement 3. Nrp-2 is enriched in the Golgi apparatus in neural tissue. Figure supplement 3—source data 1. Raw, unedited blot from Figure 3*—figure supplement 3B*. Figure supplement 3—source data 2. Raw, unedited blot from Figure 3*—figure supplement 3B*. Figure supplement 3—source data 3. Raw, labeled blot from Figure 3*—figure supplement 3B*. Figure supplement 3—source data 4. Raw, labeled blot from Figure 3*—figure supplement 3B*. Figure supplement 4. Nrp-2 palmitoyl acceptor cysteines are not required for Nrp-2/PlexA3 association but are required for proper Nrp-2 homooligomerization. Figure supplement 4—source data 1. Raw, unedited blot from Figure 3*—figure supplement 4A*. Figure supplement 4—source data 2. Raw, unedited blot from Figure 3*—figure supplement 4A*. Figure supplement 4—source data 3. Raw, unedited blot from Figure 3*—figure supplement 4A*. Figure supplement 4—source data 4. Raw, unedited blot from Figure 3*—figure supplement 4A*. Figure supplement 4—source data 5. Raw, labeled blot from Figure 3*—figure supplement 4A*. Figure supplement 4—source data 6. Raw, labeled blot from Figure 3*—figure supplement 4A*. Figure supplement 4—source data 7. Raw, labeled blot from Figure 3*—figure supplement 4A*. Figure supplement 4—source data 8. Raw, labeled blot from Figure 3*—figure supplement 4A*. Figure supplement 4—source data 9. Raw data for Figure 3*—figure supplement 4B*. Figure supplement 4—source data 10. Raw, unedited blot from Figure 3*—figure supplement 4D*. Figure supplement 4—source data 11. Raw, unedited blot from Figure 3*—figure supplement 4D*. Figure supplement 4—source data 12. Raw, unedited blot from Figure 3*—figure supplement 4D*. Figure supplement 4—source data 13. Raw, labeled blot from Figure 3*—figure supplement 4D*. Figure supplement 4—source data 14. Raw, labeled blot from Figure 3*—figure supplement 4D*. Figure supplement 4—source data 15. Raw, labeled blot from Figure 3*—figure supplement 4D*. Figure supplement 4—source data 16. Raw data for Figure 3*—figure supplement 4E*. Figure supplement 4—source data 17. Raw, unedited blot from Figure 3*—figure supplement 4F*. Figure supplement 4—source data 18. Raw, unedited blot from Figure 3*—figure supplement 4F*. Figure supplement 4—source data 19. Raw, unedited blot from Figure 3*—figure supplement 4F*. Figure supplement 4—source data 20. Raw, labeled blot from Figure 3*—figure supplement 4F*. Figure supplement 4—source data 21. Raw, labeled blot from Figure 3*—figure supplement 4F*. Figure supplement 4—source data 22. Raw, labeled blot from Figure 3*—figure supplement 4F*. Figure supplement 4—source data 23. Raw data for Figure 3*—figure supplement 4G*.

Second, we examined Nrp-2 cell surface localization in cortical neurons in culture. We transfected cortical neurons with plasmids expressing pHluorin-tagged wild-type Nrp-2 or various CS mutants and carried out live imaging. The pH-sensitive GFP variant pHluorin fluoresces robustly at neutral pH, and so the tagged protein is readily visualized when localized at the cell surface. These live imaging assessments of surface protein localization showed that Nrp-2^wild-type^ (Figure 3—figure supplement 2A; Video 1 also shows time-lapse imaging of another neuron expressing pHluorin-tagged Nrp-2^wild-type^) and Nrp-2^C922S/C923S^ (Figure 3—figure supplement 2C) exhibit a prominent punctate localization along the plasma membrane, consistent with their distribution in COS-7 cells. In contrast, Nrp-2^TCS^ apparently forms larger protein clusters (Figure 3— figure supplement 2B), and Nrp-2^Full^ ^CS^ is profoundly mislocalized, displaying a diffuse distribution over all membranes (Figure 3—figure supplement 2D). These pHluorin-tagged Nrp-2 plasmids were also tested with live imaging in Neuro2A cells, with similar results (data not shown).

Given the surface mislocalization of palmitoylation-deficient Nrp-2, we next asked how palmitoylation might affect Nrp-2 surface protein localization. Since the Golgi apparatus is the main intracellular compartment where palmitoylation enzymes are localized and function (Ohno et al., 2006; Rocks et al., 2010), we asked whether Nrp-2 localizes to Golgi membranes. We stained primary cortical neurons in culture with antibodies directed against Nrp-2 and the *cis*-Golgi marker Golgi matrix protein 130 (GM130), which showed that endogenous Nrp-2 is robustly associated with somatic Golgi and dendritic Golgi cisternae known as Golgi outposts (Figure 3—figure supplement 3A). Further, Golgi isolation experiments from mouse brain revealed Nrp-2 enrichment in the GM130-positive Golgi fraction (Figure 3—figure supplement 3B).

Next, we examined whether palmitoylated Nrp-2 cysteine residues are required for the association between Nrp-2 and the Golgi apparatus by performing immunofluorescence on *Nrp-2^-/-^* cortical neurons transfected with either wild-type Nrp-2 or various Nrp-2 CS mutants. These experiments show that wild-type Nrp-2 and Nrp-2^C922S/C923S^ proteins display robust association with Golgi membranes, whereas the Nrp-2^TCS^ and Nrp-2^Full^ ^CS^ mutant proteins are significantly deficient in their association with the Golgi apparatus (Figures 3C and 3D). The aberrant association of TCS and Full CS Nrp-2 proteins with the Golgi apparatus could have major effects on their trafficking and plasma membrane insertion.

To investigate the role of palmitoylation in the spatial and temporal dynamics of Nrp-2 trafficking, we focused on the role of transmembrane/juxtamembrane cysteines that have the greatest effect on Nrp-2 surface localization and Golgi association, performing fluorescence recovery after photobleaching (FRAP) in cortical neurons expressing exogenous pHluorin-tagged Nrp-2^wild-type^ or Nrp-2^TCS^. The fluorescent signal was bleached over a selected region of interest (ROI) and after photobleaching neurons were imaged for 15 minutes to visualize the recovery of fluorescence in the bleached area, which represents protein molecules either derived from nearby plasma membrane compartments or newly inserted in the cell membrane. We observed that Nrp-2^TCS^ protein displays a markedly different trafficking pattern with a higher mobile fraction compared to wild-type Nrp-2 (Figures 3E and 3F, and Supplement—Videos 1 and 2). This observation is in accordance with other studies reporting higher diffusibility and an altered intracellular localization of depalmitoylated proteins (Miura et al., 2006; Rocks et al., 2010, 2005).

Taken together, these experiments support the functional importance of transmembrane/juxtamembrane Nrp-2 cysteines in regulating the localization and trafficking of Nrp-2 protein across subcellular compartments and the plasma membrane.

### Nrp-2 palmitoyl acceptor cysteines are dispensable for the association between Nrp-2 and PlexA3 but are required for Nrp-2 homo-oligomerization

Sema3F binds Nrp-2, which forms homo-oligomers and interacts with plexinA3 (PlexA3) to form the Sema3F holoreceptor complex (Giger et al., 1998; Janssen et al., 2012). Since cysteine residues can influence protein structure, we asked whether Nrp-2 cysteines control the ability of Nrp-2 to associate with PlexA3 or affect the ability of Nrp-2 to homo-oligomerize.

First, we performed coimmunoprecipitation experiments using myc-tagged PlexA3 and either flag-tagged wild-type Nrp-2 or flag-tagged Nrp-2^Full^ ^CS^. These experiments showed that PlexA3 binds Nrp-2^Full^ ^CS^ to a similar extent as it does to wild-type Nrp-2 (Figure 3—figure supplement 4A and 4B). Further investigation into the association between palmitoylation-deficient Nrp-2 with PlexA3 using immunofluorescence in primary cortical neurons in culture showed that exogenous Nrp-2^Full^ ^CS^ exhibits strong co-localization with endogenous PlexA3, similar to exogenously expressed wild-type Nrp-2 (Figure 3—figure supplement 4C). Thus, the membrane-spanning and cytoplasmic Nrp-2 cysteines are dispensable for the interaction between Nrp-2 and PlexA3.

Next, we addressed whether the transmembrane and cytoplasmic Nrp-2 cysteines are required for Nrp-2 homo-oligomerization. We performed coimmunoprecipitation experiments using flag-tagged wild-type Nrp-2 and pHluorin-tagged wild-type Nrp-2 or Nrp-2^Full^ ^CS^. These experiments revealed that pHluorin-Nrp-2^Full^ ^CS^ associates with flag-Nrp-2^WT^ to a significantly lesser extent than pHuorin-Nrp-2^WT^ does (Figure 3—figure supplement 4D and 4E). Moreover, coimmunoprecipitation experiments between flag-Nrp-2^WT^ and pHluorin-Nrp-2^WT^ or Nrp-2^TCS^ demonstrated that pHluorin-Nrp-2^TCS^ associates with flag-Nrp-2^WT^ to a lesser extent than pHluorin-Nrp-2^WT^ does (Figure 3— figure supplement 4F and 4G), although this difference is not as robust as it is with Nrp-2^Full^ ^CS^. These findings show that the Nrp-2 cysteine residues that we have shown to be palmitoylated and required for Nrp-2 localization are also necessary for Nrp-2 homo-oligomerization.

### Transmembrane/juxtamembrane Nrp-2 cysteines are required for Sema3F/Nrp-2-dependent dendritic spine pruning in deep layer cortical neurons in vitro and in vivo

Sema3F/Nrp-2 signaling regulates pruning of supernumerary dendritic spines on apical dendrites of deep layer cortical pyramidal neurons (Demyanenko et al., 2014; Duncan et al., 2021; Tran et al., 2009). To investigate a potential role for palmitoyl acceptor Nrp-2 cysteines in Sema3F/Nrp-2-dependent spine constraint, we employed an in vitro assay aimed at rescuing the inability of Sema3F to decrease dendritic spine density in homozygous Nrp-2 knockout (*Nrp-2^-/-^)* cortical neurons by introducing flag-tagged wild-type Nrp-2 or Nrp-2^CS^ mutant expression plasmids subcloned in a *pCIGG2-ires-EGFP* backbone vector. *Nrp-2^-/-^* primary cortical neurons were transfected with backbone vector as a control or a Nrp-2 expression plasmid. At DIV21, when dendritic spines are well developed, neurons were treated with 5 nM Sema3F-AP or 5 nM AP (control) for 6 hrs, followed by EGFP immunofluorescence to visualize neuronal morphology. The rescue ability of the various Nrp-2 plasmids was assessed using two criteria: (1) a comparison of dendritic spine density between AP-treated *Nrp-2^-/-^* neurons expressing backbone vector (control) and AP-treated neurons expressing each *Nrp-2^CS^* mutant; and (2) a comparison of dendritic spine density between Sema3F-AP-treated and control AP-treated *Nrp-2^-/-^* cortical neurons expressing a specific Nrp-2 plasmid. If a plasmid is able to rescue the *Nrp-2* knockout spine phenotype, it should: (1) constrain spine number compared to the backbone control vector in the absence of exogenous Sema3F treatment and/or (2) cause a reduction in spine number upon exogenous Sema3F-AP treatment compared to AP treatment leading to dendritic spine pruning that is similar to that imparted by *Nrp-2^WT^*.

These experiments show that *Nrp-2^WT^* and *Nrp-2^C922S/C923S^*can rescue the *Nrp-2* null dendritic spine phenotype (Figures 4A-4C). This is in accordance with our finding that the C-terminal Nrp-2 cysteines regulate Nrp-2 subcellular localization to a rather minor extent (Figures 3A-3D, Figure 3—supplement 1, Figure 3—supplement 2). In contrast, *Nrp-2^C878S^* and *Nrp-2^C887S^* are not capable of constraining spine number, whereas *Nrp-2^C885S^* and *Nrp-2^C897S^* can partially rescue the *Nrp-2* knockout dendritic spine phenotype (Figures 4A-4C). Likewise, *Nrp-2* null rescue experiments in cortical neuron cultures with pHluorin-tagged *Nrp-2^WT^* or *Nrp-2^CS^*mutant expression plasmids including the *Nrp-2^TCS^*, *Nrp-2^C922S/C923S^* and *Nrp-2^Full^ ^CS^* mutants, reveal a requirement for transmembrane/juxtamembrane cysteines, but not the two C-terminal cysteines, in Sema3F/Nrp-2-dependent dendritic spine pruning (Figure 4—supplement 1A and 1B). The relative requirements for Nrp-2 palmitoyl acceptor cysteines in Sema3F/Nrp-2-mediated dendritic spine pruning are summarized in Figure 4D.

**Figure 4.**
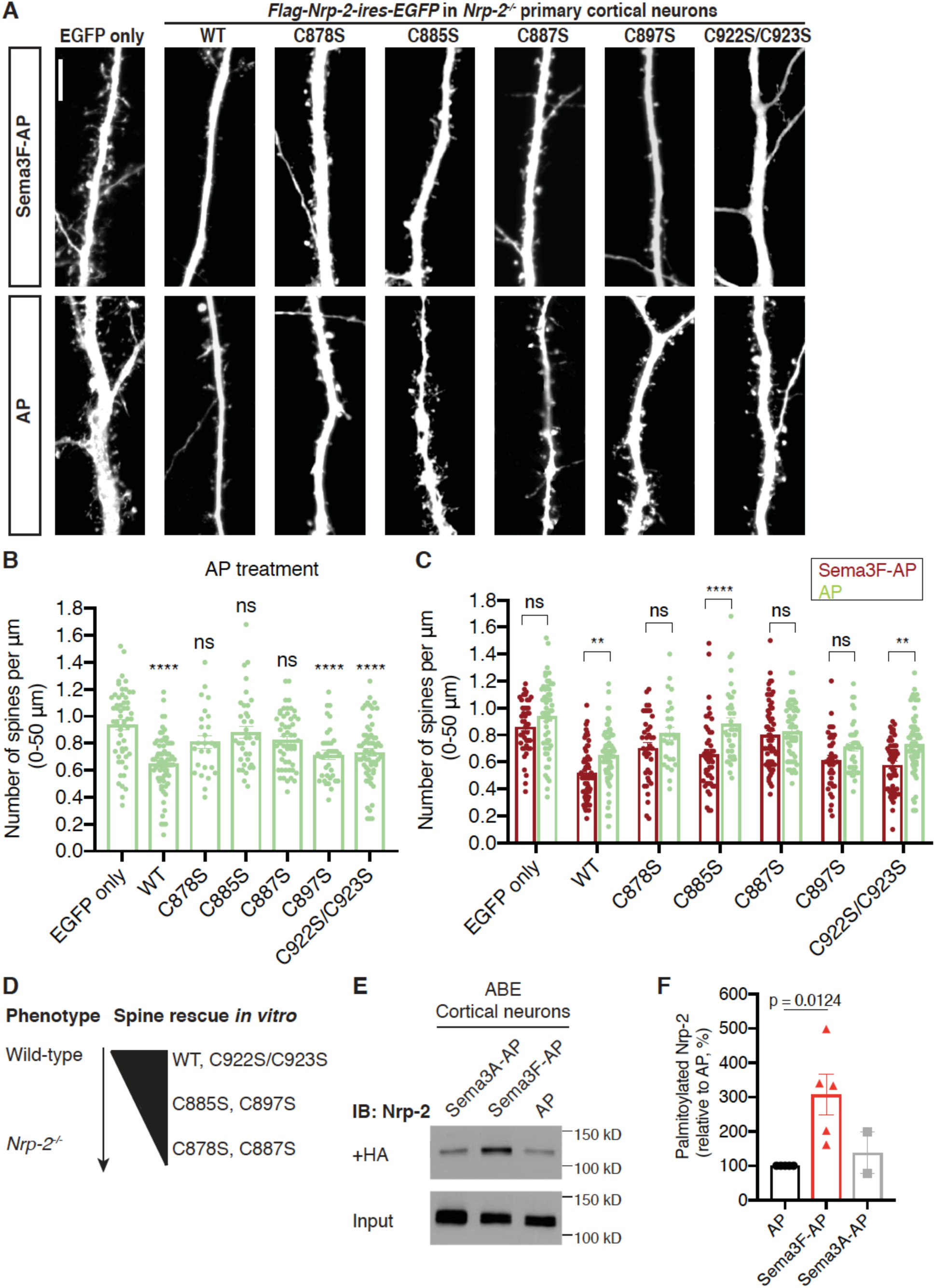
Select Nrp-2 cysteines are required for Sema3F/Nrp-2-dependent dendritic spine pruning in deep layer primary cortical neurons. **(A-D)** Rescuing in vitro the *Nrp-2^-/-^* dendritic spine phenotype to assess the role of individual Nrp-2 cysteines in Sema3F/Nrp-2-dependent dendritic spine collapse. E14.5 *Nrp-2^-/-^* primary cortical neurons were transfected with various *flag-Nrp-2-ires-EGFP* expression plasmids, including WT and CS point mutants. At DIV21 they were treated with 5 nM Sema3F-AP or 5 nM AP alone, for 6 hrs, and next subjected to EGFP immunofluorescence. (A) Representative images of neurons expressing the indicated plasmid and treated with either Sema3F-AP or AP. Images represent the 3D projection of confocal stack. Scale bar in (A), 7 µm. (**B, C**) Quantification of dendritic spines, counted along the 0-50 µm apical dendritic segment and presented as number of spines per µm. Data are plotted as mean ± SEM. The rescue ability of each Nrp-2 protein is assessed in two ways: the ability of AP-treated neurons expressing each Nrp-2 protein to constrain spines compared to AP-treated neurons expressing the backbone vector only (B) and the ability of neurons expressing each protein to respond to Sema3F-AP compared to AP (C). Graph in (B): One-way ANOVA (p < 0.0001) followed by Bonferroni’s test for multiple comparisons (compared to EGFP only: WT, p < 0.0001; C878S, p = 0.1215; C885S, p > 0.9999; C887S, p = 0.0628; C897S, p < 0.0001; C922S/C923S, p < 0.0001). Graph in (C): Two-way ANOVA (p < 0.0001) followed by Sidak’s test for multiple comparisons (Sema3F-AP vs AP: EGFP only, p = 0.4885; WT, p = 0.0074; C878S, p = 0.3555; C885S, p < 0.0001; C887S, p = 0.9985; C897S, p = 0.3308; C922S/C923S, p = 0.0017). (D) Graphic summary of the ability of tested Nrp-2 proteins to rescue the *Nrp-2^-/-^* dendritic spine density phenotype. Wild-type (WT) and C922S/C923S Nrp-2 proteins constrain dendritic spines, whereas certain CS mutants have either compromised (C885S, C897S) or abolished (C878S, C887S) rescue ability. Sema3F-AP treated: EGFP only, n = 44; WT, n = 63; C878S, n = 39; C885S, n = 46; C887S, n = 60; C897S, n = 38; C922S/C923S, n = 53. AP treated: EGFP only, n = 55; WT, n = 67; C878S, n = 26; C885S, n = 36; C887S, n = 53; C897S, n = 36; C922S/C923S, n = 64; where n is the number of neurons analyzed for each condition. (**E, F**) Sema3F enhances endogenous Nrp-2 palmitoylation in cortical neuron cultures. ABE on DIV18 primary cortical neurons treated with either 5 nM AP or 5 nM Sema3F-AP or 5 nM Sema3A-AP. (**E**) Immunoblots show palmitoylated (+HA) and input samples. –HA samples were processed in parallel with no evidence of non-specific signal (not shown). (**F**) Quantification of palmitoylated Nrp-2 levels, calculated as the ratio of +HA to the respective input. Cumulative data are plotted in scatter dot plots including the mean ± SEM. Sema3F-AP and Sema3A-AP conditions are expressed relative to AP control (set at 100%). One-way ANOVA followed by Dunnett’s test for multiple comparisons (Sema3A-AP vs AP, p = 0.8531); AP, Sema3F-AP, n = 5 independent experiments; Sema3A-AP, n = 2 independent experiments). The online version of this article includes the following source data and figure supplement(s) for figure 4: Source data 1. Raw data for Figure 4B and *4C*. Source data 2. Raw, unedited blot from Figure 4E. Source data 3. Raw, labeled blot from Figure 4E. Source data 4. Raw data for Figure 4F. Figure supplement 1. Sema3F/Nrp-2-dependent dendritic spine pruning in deep layer cortical neurons depends on distinct Nrp-2 cysteine clusters. Figure supplement 1—source data 1. Raw data for Figure 4*—figure supplement 1B*.

To investigate whether Sema3F effects on cortical neuron dendritic spines are associated with Nrp-2 palmitoylation, we asked whether Sema3F bath application impacts Nrp-2 palmitoylation. We treated wild-type cortical neurons with 5 nM Sema3F-AP, Sema3A-AP or AP and assessed palmitoylation of endogenous Nrp-2. Interestingly, Sema3F application enhances Nrp-2 palmitoylation in deep layer primary cortical neurons in culture (Figures 4E and 4F). This shows that Sema3F can upregulate Nrp-2 palmitoylation, presumably at nearby subcellular domains and within dendritic spines, and it further suggests that Sema3F effects on cortical neurons are mediated by Nrp-2 palmitoylation.

Next, we addressed whether Nrp-2 cysteines C878, C885 and C887, which are required for rescuing the Sema3F/Nrp-2-dependent dendritic spine phenotype in *Nrp-2* knockout cortical neurons in vitro, are also required for Sema3F/Nrp-2-dependent dendritic spine pruning in vivo. We examined the ability of *Nrp-2^TCS^* to constrain excess dendritic spines resulting from Nrp-2 deletion in the mouse cortex using an *in utero* electroporation (IUE) rescue paradigm. To overcome breeding challenges, including reduced viability encountered in the *Nrp-2^-/-^* mouse line, we crossed homozygous *Nrp-2* null mice with homozygous *Nrp-2 floxed* (*Nrp-2^F/F^*) mice and targeted layer V cortical pyramidal neurons using IUE in *Nrp-2^F/-^*embryos of pregnant females at E13.5. To assess the feasibility of this approach, we first carried out two control IUE experiments: one with an EGFP-expressing plasmid to visualize neuronal morphology (control) and a second using a combination of three plasmids: an *EGFP*-expressing plasmid, a *Cre*-expressing plasmid to excise the floxed *Nrp-2* allele, and a *LSL-tdTomato*-expressing plasmid that serves as a reporter of Cre activity. In neurons electroporated with all three plasmids, we expected Cre to excise the floxed *Nrp-2* allele rendering neurons homozygous *Nrp-2* knockout (*Nrp-2^-/-^*) and also to turn on *tdTomato* expression (Figure 5A). These experiments revealed increased spine density in neurons lacking Nrp-2 compared to control (Figures 5B and 5C), confirming that this approach is a useful assay for rescue analysis of the *Nrp-2^-/-^*-associated dendritic spine phenotype. Next, to assess the rescue ability Nrp-2^wild-type^ and Nrp-2^TCS^ proteins in vivo, we used two additional IUE conditions: one with the plasmid combination [*Cre* + *LSL-tdTomato* + *flag-Nrp-2^wild-type^-ires-EGFP*] and one with the plasmid combination [*Cre* + *LSL-tdTomato* + *flag-Nrp-2^TCS^-ires-EGFP*]. Neurons electroporated with the combination expressing wild-type Nrp-2 showed significantly fewer spines compared to *Nrp-2^-/-^* neurons that do not express Nrp-2 (electroporated with [*Cre* + *LSL-tdTomato* + *EGFP*]). However, neurons electroporated with the combination of plasmids that includes *Nrp-2^TCS^* exhibited a spine density similar to that of *Nrp-2^-/-^* neurons (Figures 5B and 5C).

**Figure 5.**
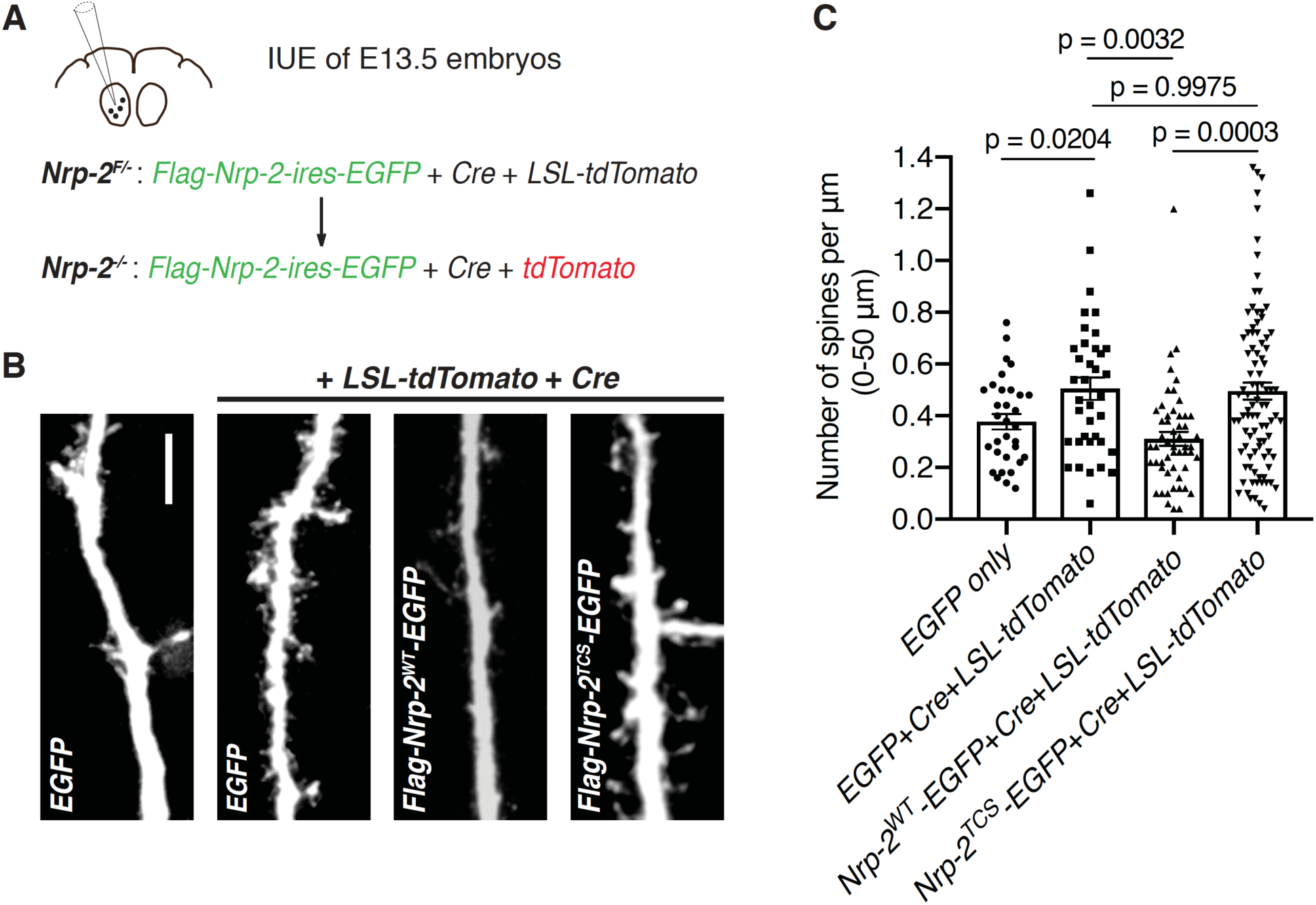
Transmembrane/juxtamembrane Nrp-2 cysteines are required in vivo for Nrp-2-dependent dendritic spine pruning in deep layer cortical pyramidal neurons. **(A-C)** Rescuing in vivo the *Nrp-2^-/-^* dendritic spine phenotype to assess the role of membrane-associated Nrp-2 cysteines in Sema3F/Nrp-2-dependent dendritic spine retraction in layer V cortical pyramidal neurons. (A) Schematic representation of the in utero electroporation (IUE) experimental approach. E13.5 *Nrp-2^F/-^* embryos were electroporated in utero either with EGFP or with [*EGFP* + *Cre* + *loxP-STOP-loxP-tdTomato* (*LSL-tdTomato*)] to excise the floxed Nrp-2 allele and render individual neurons *Nrp-2^-/-^*. Floxed tdTomato (*LSL-tdTomato*) serves as a reporter of Cre activity. These two electroporations serve as controls to assess the utility of this approach in reproducing the *Nrp-2^-/-^* spine density phenotype. Next, embryos were electroporated with either [*flag-Nrp-2^WT^-ires-EGFP* + *Cre* + *LSL-tdTomato*] or [*flag-Nrp-2^TCS^-ires-EGFP* + *Cre* + *LSL-tdTomato*] to assess the ability of WT or TCS Nrp-2 to constrain supernumerary dendritic spines resulting from Nrp-2 deletion. All mice were analyzed on postnatal day 28 (P28). (B) Representative images of apical dendrites of electroporated layer V pyramidal neurons (3D projection of confocal stack) expressing the indicated combinations of plasmids. Scale bar, 7 µm. (C) Quantification of dendritic spines, counted along the proximal 50 µm (relative to the cell body) of the apical dendrite and presented as number of spines per µm. Cumulative data from several independent IUE experiments (at least 3 mouse brains analyzed per scheme of injected plasmids) are plotted in scatter dot plots with mean ± SEM. One-way ANOVA (p = 0.0001) followed by Tukey’s test for multiple comparisons (*EGFP only*, n = 32; [*EGFP* + *Cre* + *LSL-tdTomato*], n = 37; [*flag-Nrp-2^WT^-ires-EGFP* + *Cre* + *LSL-tdTomato*], n = 53; [*flag-Nrp-2^TCS^-ires-EGFP* + *Cre* + *LSL-tdTomato*], n = 92; where n is the number of neurons analyzed). The online version of this article includes the following source data for figure 5: Source data 1. Raw data for Figure 5C.

Taken together, both in vitro and in vivo rescue experiments of the *Nrp-2^-/-^* dendritic spine density phenotype reveal an essential role for the transmembrane and membrane-proximal Nrp-2 cysteines in mediating Sema3F/Nrp-2-dependent spine pruning of layer V cortical pyramidal neurons.

### Palmitoyltransferase DHHC15 palmitoylates Nrp-2 and is required for Sema3F/Nrp-2-dependent dendritic spine pruning in primary cortical neurons

To identify the DHHC enzymes that palmitoylate Nrp-2, we carried out a Nrp-2 palmitoylation screen in 293T cells by co-expressing exogenous wild-type Nrp-2 and each of the 23 DHHCs (Fukata et al., 2004) and assessing palmitoylation with the ABE assay. Of the 23 DHHCs, DHHC15 was among the three DHHCs that most strongly enhanced Nrp-2 palmitoylation (data not shown). DHHC15 is known to palmitoylate PSD-95 (Fukata et al., 2004), is strongly expressed in the mouse cerebral cortex (Mejias et al., 2021) and is enriched in Golgi membranes (Ohno et al., 2006). Therefore, we examined the role of DHHC15 in Nrp-2 palmitoylation and function in cortical neurons.

We employed a loss-of-function approach using a *DHHC15* null mutant mouse line (Mejias et al., 2021), exploring whether Nrp-2 is a DHHC15 substrate by performing ABE in wild-type and homozygous *DHHC15* knockout (*DHHC15^-/-^*) primary cortical neurons. In *DHHC15^-/-^* cortical neurons overall Nrp-2 palmitoylation is substantially reduced (∼50%) compared to that in wild-type neurons (Figures 6A and 6B). Unlike Nrp-2, Nrp-1 palmitoylation in *DHHC15^-/-^* neurons is comparable to that observed in wild-type neurons (Figures 6A and 6C). To test for DHHC-substrate specificity, we assessed Nrp-2 palmitoylation in another previously characterized DHHC null mouse line, *DHHC8^-/-^*(Mukai et al., 2015, 2008, 2004), since DHHC8 did not enhance Nrp-2 palmitoylation in our screen in 293T cells (data not shown). Nrp-2 palmitoylation levels in *DHHC8^-/-^* cortical neurons are very close to those observed in wild-type cortical neurons (Figure 6—figure supplement 1A and 1B) and, therefore, Nrp-2 does not appear to be a palmitoyl substrate of DHHC8 in these neurons.

**Figure 6.**
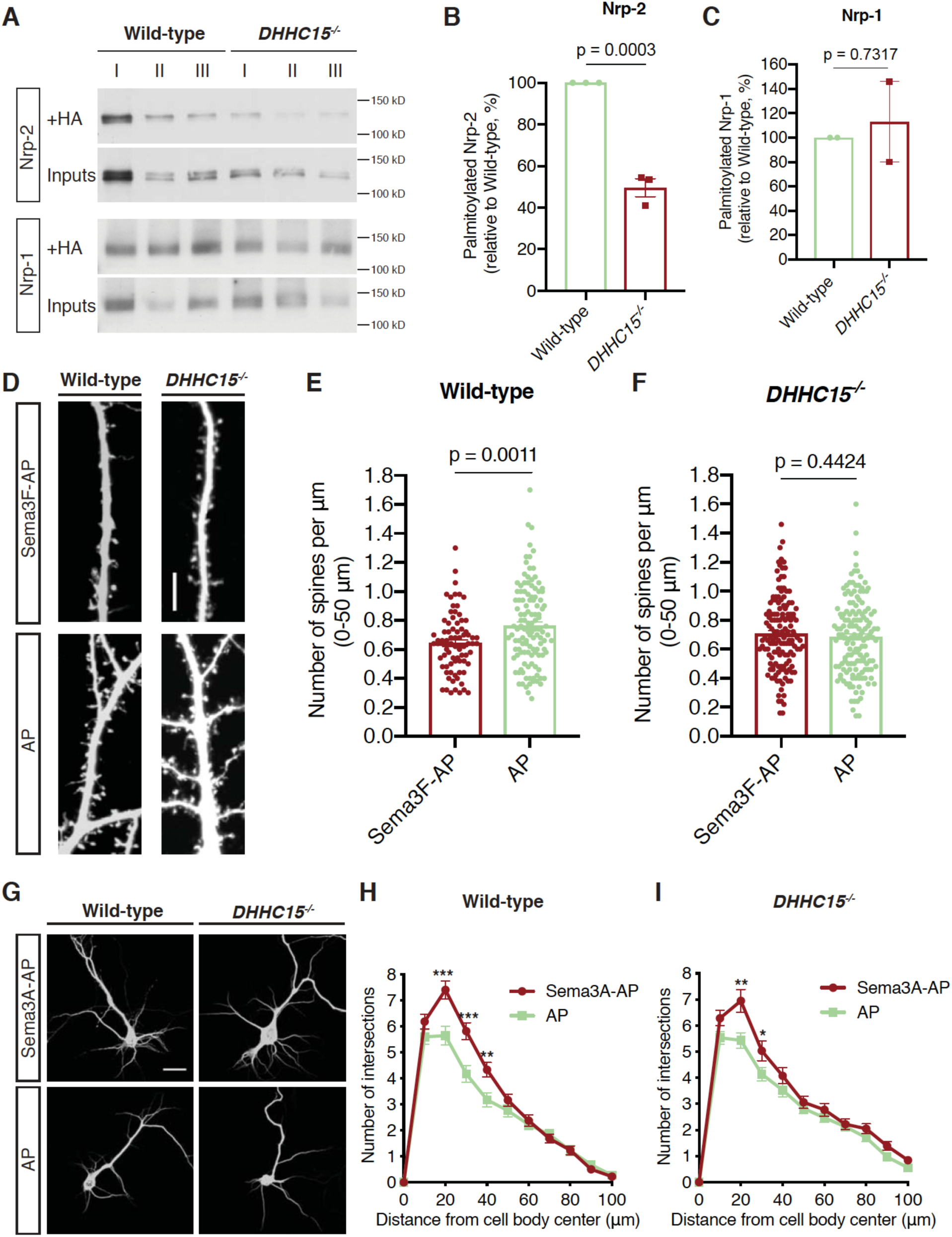
DHHC15 regulates Nrp-2 palmitoylation and function, but not Nrp-1, in primary cortical neurons. **(A-C)** ABE performed on E14.5 DIV12 wild-type C57BL/6 or *DHHC15^-/-^* primary cortical neurons. (A) Representative immunoblots of palmitoylated (+HA) and input samples for Nrp-2 and Nrp-1. (B, C) Quantification of palmitoylated Nrp-2 (B) or Nrp-1 (C), calculated as the ratio of palmitoylated protein (+HA) to the respective input; this ratio for wild-type (WT) neurons is set at 100% and the ratio for *DHHC15*^-/-^ neurons is expressed as a percentage of WT. In each independent experiment, three different samples (I, II, III; replicates), for each genotype, were processed in parallel and averaged. Nrp-2, n = 3 independent experiments; Nrp-1, n = 2 independent experiments. Pooled data are plotted in scatter dot plots with mean ± SEM. Two-tailed t test; Nrp-2 mean: WT = 100%, *DHHC15^-/-^* = 49.64%; Nrp-1 mean: WT = 100%, *DHHC15^-/-^* = 113%. **(D-F)** Assessment of the ability of wild-type C57BL/6 or *DHHC15^-/-^* primary cortical neurons to respond to Sema3F by excess dendritic spine pruning. Neurons were transfected with an EGFP-expressing plasmid to visualize neuronal morphology and at DIV21 they were treated with 5 nM Sema3F-AP or 5 nM AP (control) for 6 hrs. Spines are quantified along the cell body-proximal 50 µm of the apical dendrite. (D) Representative images are shown for wild-type and *DHHC15^-/-^* neurons. Scale bar, 7 µm. (E, F) Quantification of dendritic spines, counted along the proximal 50 µm (from cell body) on the largest dendrite, presented as number of spines per µm and plotted in scatter dot plots including mean ± SEM (3 independent experiments). Note that wild-type neurons exhibit significant Sema3F-induced spine retraction, whereas *DHHC15^-/-^*neurons invariably display no response to Sema3F-AP (compared to AP). Wild-type: Sema3F-AP, n = 78; AP, n = 119. *DHHC15^-/-^*: Sema3F-AP, n = 144; AP, n = 147, where n is the number of neurons analyzed for each condition. Two-tailed t test. **(G-I)** Assessment of the ability of wild-type C57BL/6 or *DHHC15^-/-^* primary cortical neurons to respond to Sema3A by elaborating their dendritic tree. E14.5 primary cortical neurons treated at DIV12 with 5 nM Sema3A-AP or 5 nM AP (control) for 6 hrs and subjected to MAP2 immunofluorescence. (G) Representative images are shown for each genotype and represent the 3D projection of confocal stack. Scale bar, 20 µm. (H, I) Quantitative assessment of dendritic arborization with Sholl analysis, presented as number of intersections at various distances from the center of the cell body and plotted as mean ± SEM for each distance (3 independent experiments). Both wild-type and *DHHC15^-/-^* cortical neurons exhibit a more elaborate perisomatic dendritic arbor following Sema3A-AP treatment compared to AP (control) treatment. Wild-type: Sema3A-AP, n = 72; AP, n = 62. *DHHC15^-/-^*: Sema3A-AP, n = 77; AP, n = 110, where n is the number of neurons analyzed. Multiple t tests; (H), *** p < 0.001, **p = 0.0043; (I), **p = 0.003, *p = 0.046. The online version of this article includes the following source data and figure supplement(s) for figure 6: **Source data 1.** Raw, unedited blot from Figure 6A. **Source data 2.** Raw, unedited blot from Figure 6A. **Source data 3.** Raw, labeled blot from Figure 6A. **Source data 4.** Raw, labeled blot from Figure 6A. **Source data 5.** Raw data for Figure 6B. **Source data 6.** Raw data for Figure 6C. **Source data 7.** Raw data for Figure 6E. **Source data 8.** Raw data for Figure 6F. **Source data 9.** Raw data for Figure 6H. **Source data 10.** Raw data for Figure 6I. **Figure supplement 1.** Nrp-2 is not a substrate for the palmitoyl acyltransferase DHHC8 in deep layer primary cortical neurons. **Figure supplement 1—source data 1.** Raw, unedited blot from Figure 6***—figure supplement 1A***. **Figure supplement 1—source data 2.** Raw, labeled blot from Figure 6***—figure supplement 1A***. **Figure supplement 1—source data 3.** Raw data for Figure 6***—figure supplement 1B***. **Figure supplement 1—source data 4.** Raw data for Figure 6***—figure supplement 1D***.

Next, we asked whether DHHC15 plays a role in Sema3F/Nrp-2-dependent dendritic spine pruning. Wild-type and *DHHC15^-/-^*deep layer primary cortical neurons were cultured, transfected with an EGFP-expressing plasmid, and at DIV21 they were treated with 5 nM Sema3F-AP or 5 nM AP for 6 hrs. They were then subjected to EGFP immunofluorescence, imaged and spine density was quantified along the largest dendrite. Unlike wild-type cortical neurons, which displayed a significant reduction in spine density following treatment with Sema3F-AP compared to AP (Figures 6D and 6E), *DHHC15^-/-^* neurons failed to exhibit apical dendritic spine pruning following Sema3F-AP treatment (Figures 6D and 6F).

To address the specificity of these spine morphology effects, we tested the ability of cortical neurons lacking DHHC8, which plays important roles in the nervous system (Mukai et al., 2015, 2008), to exhibit spine pruning in response to Sema3F. *DHHC8^-/-^* cortical neuron cultures were transfected with an EGFP expression plasmid, and at DIV21 they were treated with Sema3F-AP or AP for 6 hrs followed by EGFP immunofluorescence. *DHHC8^-/-^* cortical neurons responded to Sema3F with significant spine retraction (Figure 6—figure supplement 1C and 1D), similar to wild-type cortical neurons.

Since we observed no effect of *DHHC15* loss-of-function on Nrp-1 palmitoylation, we investigated whether DHHC15 is required for Sema3A/Nrp-1-dependent basal dendritic elaboration (Gu et al., 2003). Wild-type and *DHHC15^-/-^*cortical neuron cultures were treated at DIV12 with 5 nM Sema3A-AP or 5 nM AP for 6 hrs. Next, they were subjected to immunofluorescence with an antibody directed against the dendritic marker microtubule-associated protein 2 (MAP2) to visualize dendritic trees and subjected to Sholl analysis. Wild-type neurons exhibited enhanced elaboration of perisomatic dendrites in response to Sema3A-AP, compared to the AP control (Figures 6G and 6H), in line with the effect of Sema3A/Nrp-1 signaling on basal dendrite elaboration of deep layer cortical pyramidal neurons in the mouse brain (Gu et al., 2003). Likewise, *DHHC15^-/-^* cortical neurons responded to Sema3A with enhanced elaboration of their basal dendrite arbors following Sema3A-AP treatment as compared to the AP control (Figures 6G and 6I).

These biochemical and phenotypic experiments in vitro reveal that DHHC15 plays essential roles in Nrp-2 palmitoylation and Sema3F/Nrp-2-dependent dendritic spine pruning, and that DHHC15 is dispensable for Nrp-1 palmitoylation or Sema3A/Nrp-1-dependent basal dendritic elaboration in primary deep layer cortical neurons. These results underscore the exquisite specificity among palmitoyl acyltransferases for their neuronal substrates.

### DHHC15 controls Nrp-2-dependent dendritic spine pruning in deep layer cortical pyramidal neurons in vivo

Since we observe that DHHC15 is required for Nrp-2 palmitoylation and Sema3F/Nrp-2-dependent dendritic spine pruning of deep layer cortical neurons in culture, we asked whether DHHC15 is also required for Nrp-2-mediated functions in cortical pyramidal neurons in the mouse neocortex. We examined dendritic spine density on the apical dendrite of layer V cortical pyramidal neurons in *DHHC15^-/-^* and wild-type mice using two different labeling techniques: a genetic labeling approach using the *Thy1-EGFP-m* mouse line that labels layer V cortical pyramidal neurons and the Golgi staining technique for sparse neuronal labeling. Our analysis of apical dendritic spines in wild-type and *DHHC15^-/-^* layer V cortical pyramidal neurons labeled with the *Thy1-EGFP-m* line shows that *DHHC15^-/-^* dendrites harbor more dendritic spines compared to wild-type neurons (Figures 7A and 7B), phenocopying previously observed effects of *Nrp-2* loss-of-function in this same neuronal population in vivo (Tran et al., 2009). Golgi-stained brains yielded similar results (Figure 7—figure supplement 1A and 1B). Next, given the lack of an effect of DHHC15 in Nrp-1 palmitoylation and function, we assessed the Sema3A/Nrp-1-dependent basal dendritic arbor elaboration in Golgi-stained wild-type and *DHHC15^-/-^* brains and observed no difference in dendritic arbor complexity between wild-type and *DHHC15^-/-^* deep layer cortical pyramidal neurons (Figure 7—figure supplement 1C and 1D).

**Figure 7.**
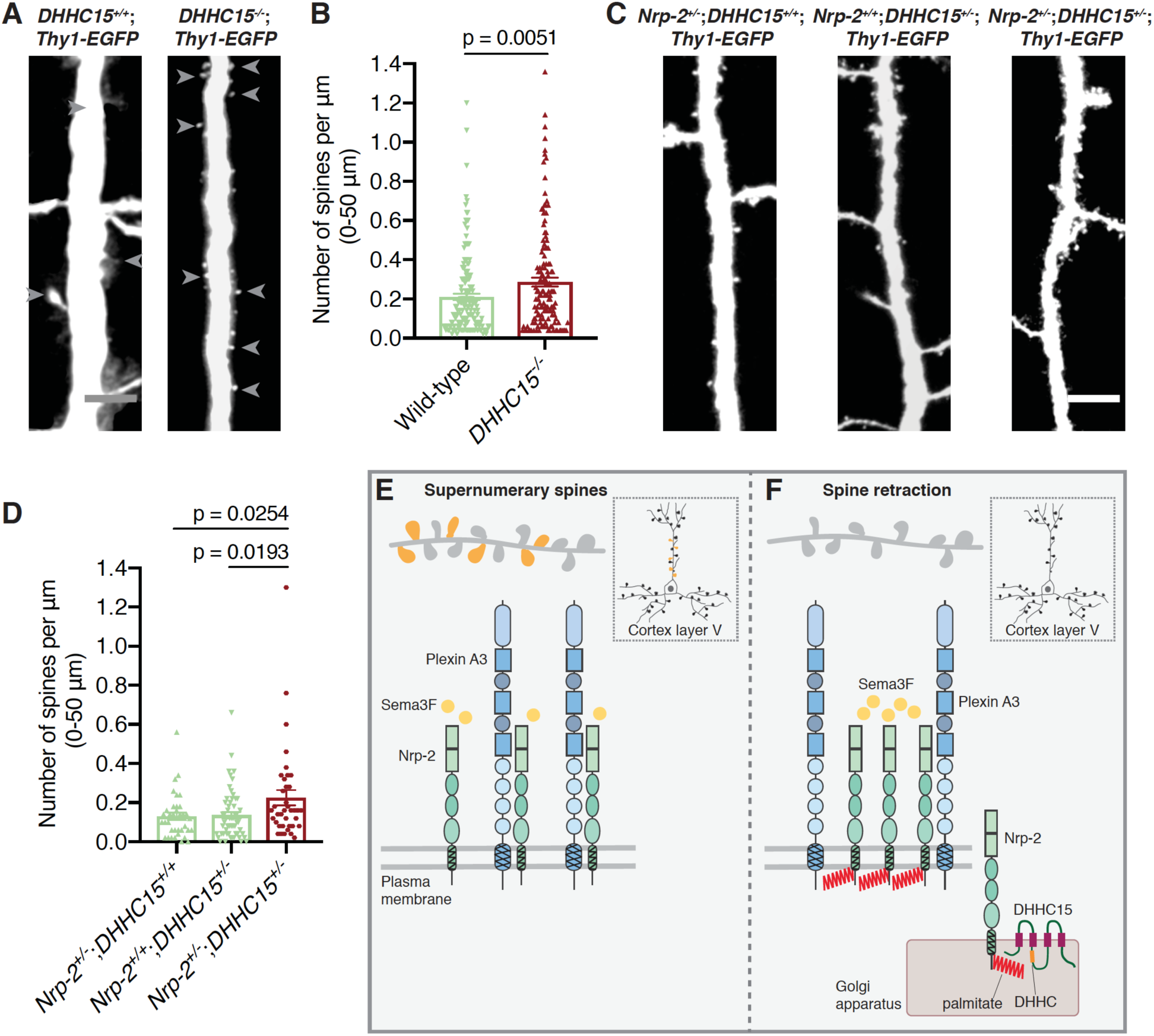
Palmitoyl acyltransferase DHHC15 and Nrp-2 interact in vivo during Nrp-2-dependent pruning of excess dendritic spines in layer V cortical pyramidal neurons. (**A, B**) Assessment of dendritic spine density on the apical dendrite of layer V cortical pyramidal neurons on postnatal day 28 (P28) *DHHC15^+/+^*;*Thy1-EGFP* and *DHHC15^-/-^*;*Thy1-EGFP* brains. (A) Panels show representative images of the apical dendrite of Thy1-EGFP-labeled layer V cortical pyramidal neurons for each genotype following EGFP immunofluorescence. Scale bar, 7 µm. (B) Quantification of dendritic spines, counted along the proximal 50 µm (relative to the cell body) of the apical dendrite, presented as number of spines per µm and plotted in scatter dot plots including mean ± SEM (3 pairs of mice analyzed). *DHHC15^+/+^*;*Thy1-EGFP*: n = 161, *DHHC15^-/-^*;*Thy1-EGFP*: n = 139, where n is the number of neurons analyzed; Two-tailed t test. (**C, D**) Genetic interaction analysis in vivo between Nrp-2 and DHHC15 for the *Nrp-2* null dendritic spine density phenotype in layer V cortical pyramidal neurons. Littermates of three different genotypes were analyzed on P33: the single heterozygotes *Nrp-2^+/-^*;*DHHC15^+/+^*;*Thy1-EGFP* and *Nrp-2^+/+^*;*DHHC15^+/-^*;*Thy1-EGFP* and the transheterozygote *Nrp-2^+/-^*;*DHHC15^+/-^*;*Thy1-EGFP*. (C) Panels show representative images of apical dendrites of Thy1-EGFP-labeled layer V cortical pyramidal neurons harboring the indicated genotype, visualized with EGFP immunofluorescence. Scale bar, 7 µm. (D) Quantification of the genetic interaction experiment between Nrp-2 and DHHC15. Dendritic spines were counted along the 0-50 µm apical dendritic segment, presented as number of spines per µm^2^ and plotted in scatter dot plots including mean ± SEM (two brains, 4 cerebral hemispheres, analyzed per genotype). *Nrp-2^+/-^*;*DHHC15^+/+^*;*Thy1-EGFP* (in green): n = 36, *Nrp-2^+/+^*;*DHHC15^+/-^*;*Thy1-EGFP* (in green): n = 57, *Nrp-2^+/-^*;*DHHC15^+/-^*;*Thy1-EGFP* (in red): n = 38, where n is the number of neurons analyzed. One-way ANOVA followed by Dunnett’s test for multiple comparisons. (E, F) Schematic illustration of the effects of Nrp-2 palmitoylation on Nrp-2 subcellular localization and Sema3F/Nrp-2-dependent dendritic spine pruning in cortical neurons. (E) Unpalmitoylated Nrp-2 is diffusely distributed over the entire plasma membrane and unable to constrain excess dendritic spines along the apical dendrite of layer V cortical pyramidal neurons (inset depicts a pyramidal neuron of the cerebral cortex). (F) Palmitoylation on transmembrane/juxtamembrane Nrp-2 cysteines, mediated in part by DHHC15 in the Golgi apparatus and enhanced by Sema3F, enables Nrp-2 clustering on distinct plasma membrane domains and Sema3F/Nrp-2-dependent pruning of supernumerary dendritic spines on the apical dendrite of layer V pyramidal neurons of the cerebral cortex. The online version of this article includes the following source data and figure supplement(s) for figure 7: Source data 1. Raw data for Figure 7B. Source data 2. Raw data for Figure 7D. Figure supplement 1. Selective effects of DHHC15 on Nrp-2-dependent developmental processes in the mouse brain. Figure supplement 1—source data 1. Raw data for Figure 7*—figure supplement 1B*. Figure supplement 1—source data 2. Raw data for Figure 7*—figure supplement 1D*.

Finally, we examined whether DHHC15 and Nrp-2 exert their effects in the same signaling pathway in layer V cortical pyramidal neurons in vivo, by performing a genetic interaction experiment. We analyzed mice of three genotypes: (1) *Nrp-2^+/-^*;*DHHC15^+/+^*;*Thy1-EGFP,* (2) *Nrp-2^+/+^*;*DHHC15^+/-^*;*Thy1-EGFP* and (3) *Nrp-2^+/-^*;*DHHC15^+/-^*;*Thy1-EGFP*. We found that transheterozygous *Nrp-2* and *DHHC15* mutant mice display a significantly higher density of dendritic spines on the apical dendrite of deep layer cortical pyramidal neurons compared to single *Nrp-2* or *DHHC15* heterozygotes (Figures 7C and 7D). These observations support the hypothesis that Nrp-2 and DHHC15 function in the same signaling pathway in vivo.

We also examined whether DHHC15 contributes to Nrp-2-dependent development and guidance of fiber tracts, including the anterior commissure and select cranial nerves (Giger et al., 2000). Analysis of wild-type and *DHHC15^-/-^* brains using neurofilament staining revealed an intact anterior commissure in *DHHC15* deficient mice (Figure 7—figure supplement 1E). Likewise, the examination of *DHHC15^-/-^* embryos with whole-mount neurofilament staining revealed normal development and guidance of cranial nerves (data not shown).

Taken together, our in vitro and in vivo data suggest a critical role for select palmitoylated Nrp-2 cysteines in Nrp-2 subcellular localization and function in cortical neurons. Further, they reveal that palmitoyl acyltransferase-substrate specificity drives diversification of Nrp-2 and Nrp-1 palmitoyl substrates to control deep layer cortical pyramidal neuron morphology and, presumably, neural circuit assembly.

## DISCUSSION

We show here that Nrp-2 and Nrp-1 are novel neuronal palmitoylation substrates and that select palmitoyl acceptor Nrp-2 cysteines are critical for Nrp-2 protein subcellular localization, for plasma membrane insertion and for Sema3F/Nrp-2-dependent apical dendritic spine pruning in deep layer cortical pyramidal neurons in vitro and in vivo (see schematics, Figures 7E and 7F). Importantly, a comparative analysis between Nrp-2 and Nrp-1 shows that palmitoyl acyltransferase-substrate specificity contributes to the functional divergence and specification of Sema3F/Nrp-2 as compared to Sema3A/Nrp-1 signaling pathways in the CNS.

The discovery and functional characterization of neuropilin palmitoylation opens new avenues for investigating how these multifaceted neuronal cue receptors sculpt neuronal circuits. Despite Nrp-2 and Nrp-1 displaying similar palmitoylation patterns in their transmembrane and juxtamembrane segments, they exhibit critical differences with regard to their subcellular localization and function, and these can be partially ascribed to palmitoylation. Specifically, palmitoylation conveys upon Nrp-2 a highly clustered localization pattern on the cell surface, whereas Nrp-1 is diffusely localized on the plasma membrane regardless of its palmitoylation state. Further, our biochemical and phenotypic analyses of Sema3F/Nrp-2-dependent cortical neuron dendritic spine pruning and Sema3A/Nrp-1-dependent cortical neuron dendritic elaboration reveal that the palmitoyl acyltransferase DHHC15 is required for Sema3F/Nrp-2 function in cortical neurons, whereas it is not required for Sema3A/Nrp-1 function in this same class of neuron. These observations show that palmitoylation endows Nrp-2 and Nrp-1 with distinct functional properties that allow these receptors to exert their well-defined and distinct effects on cortical pyramidal neuron morphogenesis. This may be explained by palmitoylation events mediated by distinct palmitoyl acyltransferases and/or palmitoylation at distinct subcellular sites being differentially important for protein localization and function. For example, palmitoylation close to the plasma membrane might regulate the targeting of a protein to dendritic spines or plasma membrane microdomains, and these events may also include activity-dependent or ligand-dependent effects on protein trafficking (Brigidi et al., 2014; Noritake et al., 2009). Additional factors that may interact with palmitoylation signaling include other posttranslational modifications (Lin et al., 2009; Salaun et al., 2010), and also sorting signals and protein-protein interactions that may endow Nrp-2 and Nrp-1 with additional protein-specific localizing and functional properties.

Our finding that Nrp-2 cysteines critically control Nrp-2 localization, homomultimerization and function in cortical neurons raises the question as to whether these effects are mediated by palmitoylation *per se*, or whether they result from cysteine residue-dependent structural effects. Several lines of evidence presented here, including robust Nrp-2 palmitoylation, the effect of 2-bromopalmitate on Nrp-2 cell surface localization, the Sema3F-induced enhancement of Nrp-2 palmitoylation and the critical role of DHHC15 in Nrp-2 function, provide strong support for a palmitoylation-dependent role for these cysteines. Interestingly, cysteine palmitoylation itself has been shown to impact protein dimerization (Bhattacharyya et al., 2016). Given its reversibility, palmitoylation could give rise to bidirectional transformation between a palmitoylated state and a non-palmitoylated state that takes part in disulfide-linked dimer formation. In addition, given the critical role of neuron-glial related cell adhesion molecule (NrCAM) in Sema3F-dependent dendritic spine pruning in cortical neurons (Demyanenko et al., 2014; Duncan et al., 2021), it will be of interest to investigate the role of Nrp-2 palmitoylation in the physical and functional interactions between the Nrp-2/PlexA3 holoreceptor complex and NrCAM.

We find that the C-terminal Nrp-2 di-cysteine motif CCXXX_*_ (“_*_” denotes a stop codon) does not appear to play essential roles in Nrp-2 localization on Golgi membranes, cell surface compartmentalization or dendritic spine pruning, though minor effects of these cysteines can be observed in certain localization assays. This C-terminal motif is also present in paralemmin (Kutzleb et al., 1998) and RhoB (Adamson et al., 1992), proteins in which the upstream cysteine is palmitoylated and the downstream cysteine is prenylated (Adamson et al., 1992; Fukata and Fukata, 2010; Kutzleb et al., 1998). It is likely that the Nrp-2 C-terminal di-cysteine motif is important for other aspects of Nrp-2 function such as axon guidance or axon pruning. Further, the similar cell surface distribution patterns of Nrp-2^TCS^ and Nrp-2^Full^ ^CS^ suggest that the juxtamembrane cysteines predominate over the C-terminal cysteines in regulating Nrp-2 subcellular localization. Overall, our observations favor a model whereby functional segregation within the Nrp-2 transmembrane and cytoplasmic domains is attributable in part to distinct roles in Nrp-2 localization and function being imparted by distinct cysteine clusters, consistent with previous observations on palmitoylation effects on ionotropic receptors (Hayashi et al., 2009, 2005).

Our observation that Nrp-2 distribution includes localization in the Golgi apparatus suggests that bulk Nrp-2 palmitoylation occurs on Golgi membranes, and that palmitoylated Nrp-2 enters the post-Golgi secretory pathway to undergo anterograde trafficking toward the plasma membrane. Interestingly, a significant fraction of Nrp-2 protein is associated with Golgi membranes in dendrites of cortical neurons (Figures 3C and S4A), known as Golgi outposts (Horton et al., 2005). This suggests Nrp-2 may also be palmitoylated locally in response to stimuli including its ligand Sema3F, other signaling effectors or even neuronal activity, and that Nrp-2 is then delivered to dendritic spines and synapses in response to these cues, as has been observed for other proteins (Noritake et al., 2009). Therefore, the enhancement we observe of Nrp-2 palmitoylation by exposure to exogenous Sema3F (Figures 4E and 4F) may lead to rapid delivery of Nrp-2 to synaptic sites. Moreover, Nrp-2 and its ligand Sema3F are dynamically regulated in response to alterations in neuronal activity (Lee et al., 2012; Wang et al., 2017); in the case of Nrp-2 this may in part be controlled by rapid cycles of Nrp-2 palmitoylation-depalmitoylation in the vicinity of individual dendritic spines, a mechanism that can confer both spatial and temporal precision of protein localization and function (Fivaz and Meyer, 2003; Fukata et al., 2004). This idea is in line with studies demonstrating activity-dependent palmitoylation of synapse-associated proteins in neural tissue (Brigidi et al., 2014; Hayashi et al., 2005; Noritake et al., 2009).

The temporal and spatial organization of the palmitoylation network is largely governed by the precise spatiotemporal regulation of palmitoyl acyltransferases (Greaves and Chamberlain, 2011; Noritake et al., 2009; Ohno et al., 2006), rendering substrate specificity a major determinant of the functional specification of cues that can regulate neuronal architecture and polarity. However, the landscape of protein palmitoylation is further enriched by modulators of DHHC activity and the concurrent activity of depalmitoylating enzymes (Salaun et al., 2020; Tortosa et al., 2017; Yokoi et al., 2016). There is increasing evidence that palmitoylation deficits lead to altered neuronal excitability and aberrant neuronal phenotypes linked to neural diseases (Mansouri et al., 2005; Milnerwood et al., 2013; Mukai et al., 2015, 2008, 2004; Pinner et al., 2016; Raymond et al., 2007; Singaraja et al., 2011; Sutton et al., 2013). In particular DHHC15, shown here to play essential roles in Nrp-2 palmitoylation and function, has been implicated in X-linked intellectual disability (Mansouri et al., 2005), impairments in learning and memory (Wang et al., 2015), hyperactivity associated with a novel environment and sensitivity to psychostimulants (Mejias et al., 2021) and also dendritic outgrowth and formation of mature spines in hippocampal neurons in vitro (Shah et al., 2019). Intriguingly, Sema3F/Nrp-2 signaling has been shown to play important roles in hippocampal circuit function, and its dysregulation may cause epileptic activity in mice (Li et al., 2022; Sahay et al., 2005). Moreover, neuropilins are key players in cancer growth and progression and response to antineoplastic therapies (Napolitano and Tamagnone, 2019; Rizzolio and Tamagnone, 2011), and they also play important roles in the function of the immune system (Kumanogoh and Kikutani, 2013) and the vascular system (Gu et al., 2003; Simons et al., 2016). Our experiments identify Nrps as new members of the neuronal palmitoyl proteome, reveal new DHHC enzymatic substrates in the CNS and define DHHC enzyme-substrate specificity as a novel mechanism specifying the functional identity of neuronal substrates. Our results advance our understanding of CNS development and function and may prove invaluable for the development of targeted therapeutic approaches directed toward amelioration of neural disorders associated with aberrant function of palmitoylation signaling pathways.

## Author Contributions

Conceptualization, E.K., D.D.G. and A.L.K.; Methodology, E.K. and A.L.K.; Investigation, E.K., Q.W., R.M., and R.H.; Validation, E.K., Q.W., R.H., and A.L.K.; Formal Analysis, E.K., Q.W., and R.H; Writing – Original Draft, E.K. and A.L.K.; Writing – Review & Editing, E.K., Q.W., R.M., R.H., T.W., D.D.G., and A.L.K.; Supervision, A.L.K., D.D.G., and T.W.; Funding Acquisition, A.L.K., D.D.G., and E.K.

## Additional files

- Materials Design Analysis Reporting (MDAR) Checklist for Authors

## Acknowledgements

We thank Dontais Johnson and Sarah Mitchell for excellent technical assistance. We also thank Martin M. Riccomagno and other Kolodkin laboratory members for discussions. We thank Dr. Joseph A. Gogos for providing the *DHHC8^-/-^* mouse line and Dr. Masaki Fukata for providing DHHC expression plasmids. This work was supported by NIH grant MH100024-Project #3 (A.L.K.); The Howard Hughes Medical Institute (A.L.K., D.D.G.); and the Greek State Scholarships Foundation and the A.G. Leventis Foundation (E.K.).

## MATERIALS AND METHODS

### Animals

Animal procedures were carried out in conformity with the policies and guidelines of the Animal Care and Use Committee (ACUC) of the Johns Hopkins University (protocol # MO20M48), which are established according to the US National Research Council’s Guide to the Care and Use of Laboratory Animals and in compliance with the Animal Welfare Act and Public Health Service Policy. Mice were handled with care and every effort was made to minimize suffering. Wild-type C57BL/6 mice and Thy1-EGFP-m transgenic mice were purchased from Jackson laboratory. The *DHHC15* (or *ZDHHC15*) knockout mouse line was generated in Dr. Tao Wang’s laboratory (Johns Hopkins University) (Mejias et al., 2021). The *DHHC15* mutant mouse was crossed to pure *C57BL/6* mice to give a pure *C57BL/6* background. The *DHHC8* (or *ZDHHC8*) knockout mouse line was provided to our lab by Dr. Joseph Gogos (Columbia University, New York) and has been characterized (Mukai et al., 2008, 2004).

### Cell lines and cell line cultures

HEK (human embryonic kidney) 293T cells (ATCC, Cat no. CRL-11268 RRID: CVCL_1926, female), COS-7 (CV-1 in Origin with SV40 genes) cells (ATCC, Cat no. CRL-1651, RRID: CVCL_0224, male) and Neuroblastoma 2A (Neuro2A) cells were maintained in culture media consisting of DMEM (Dulbecco’s Modified Eagle Medium), fetal bovine serum (10% final), penicillin and streptomycin (50 U/ml) and Glutamax supplement (1x final, ThermoFisher Scientific), in a humidified incubator at 37°C with 5% CO2. Cells were plated onto 6-well dishes for biochemical experiments or onto 12-well or 24-well dishes on glass coverslips for immunofluorescence, at the desired confluency. Cells were transfected using Lipofectamine 2000 reagent (ThermoFisher Scientific), according to manufacturer’s instructions, and processed 24-48 hrs after transfection based on protein expression. These cell lines were not authenticated.

### Primary cortical neuronal cultures

Timed-pregnant female mice were either obtained from external organizations or generated in-house by plug checks. For deep layer cortical neuronal cultures the hemispheres (excluding ventral structures and the olfactory bulb) were dissected out from embryos of both sexes from timed-pregnant mice at embryonic day (E) E14.5. During dissection the dissected tissue was kept on ice-cold Leibovitz’s L-15 medium (ThermoFisher Scientific). Tissue was digested in HBSS (Hank’s balanced salt solution) containing 0.1% trypsin, in a 37°C-water bath for 15 min. Next, cortices were washed twice with HBSS containing 10% fetal bovine serum (FBS) to inactivate trypsin and dissociated in neuron growth medium containing 10% FBS by gently passing them several times through a glass Pasteur pipette. Dissociated neurons were plated onto 6-well dishes for biochemical experiments or onto 12-well or 24-well dishes on glass coverslips for immunofluorescence, at the desired confluency. Next day, medium was replaced by fresh neuron growth medium and thereafter half of the medium was changed every 1 or 2 days. Neuron growth medium consisted of Neurobasal medium (ThermoFisher Scientific, Gibco) supplemented with 2% B-27 supplement (Gibco), 2 mM Glutamax (Gibco) and 1x penicillin/streptomycin (Gibco). Neuronal cultures were maintained in a humidified incubator at 37°C with 5% CO_2_.

Dish/coverslip preparation: Round glass coverslips were treated with nitric acid overnight, next washed with ddH_2_O and ethanol and stored in 95% ethanol. The day of culture, dishes, with or without coverslips, were coated with 0.1 mg/ml poly-D-lysine (Sigma) diluted in ddH_2_O at 37°C for at least 3 hrs. Before plating, poly-D-lysine was removed and dishes were washed twice with DPBS (Dulbecco’s phosphate-buffered saline).

### Plasmids

Plasmids were generated with standard cloning techniques. Briefly, Polymerase Chain Reaction (PCR)-amplified inserts carrying the appropriate restriction sites on the 5′ and 3′ ends were digested and ligated with the backbone vector. N-terminally flag-tagged Nrp-1 and Nrp-2 expression plasmids were generated in a *pCIG2-ires-EGFP* backbone vector containing the preprotrypsin signal peptide followed by the flag epitope tag and the Nrp-1 or Nrp-2 coding sequences in frame with the above elements. N-terminally pHluorin-tagged Nrp-2 expression plasmid was generated by subcloning in frame the endogenous signal peptide of Nrp-2, the pHluorin coding sequence and the Nrp-2 coding sequence (downstream of its signal peptide) in a *pCAGGS-ires-dsRED* backbone vector. The GFP variant pHluorin displays strong fluorescence (bright green) at neutral pH, which occurs upon receptor surface localization, whereas it does not fluoresce in vesicles (pH < 6.0). As such, it provides a robust assay for cell surface protein visualization in live cells, while upon regular EGFP immunofluorescence on permeabilized cells allows total (surface and intracellular) protein visualization.

Cysteine-to-serine point mutants (CS) of Nrp-1 and Nrp-2 were generated by amplification of the wild-type protein with reverse primers harboring the desired mutation(s). All clones were fully sequenced and protein expression was assessed by western blotting in heterologous cells and immunofluorescence in cell lines and cultured neurons. The CS point mutations did not alter the molecular weight of Nrp-2 or Nrp-1, as evidenced by western blotting.

### Transfections

Transfection of COS-7 cells, 293T cells and primary cortical neurons was performed with Lipofectamine 2000 (ThermoFisher Scientific, Cat no. 11668-019) according to the manufacturer’s guide. In brief, the appropriate amounts of DNA and Lipofectamine were added in separate tubes containing Opti-MEM I medium and incubated at room temperature for 5 min. Next, diluted DNA and diluted Lipofectamine were mixed and co-incubated at room temperature for 20 min. The mix was added to cells and after ∼ 5 hrs the medium was replaced by fresh medium. Primary cortical neurons were transfected between DIV7 and DIV9. Transfection of Neuro2A cells was performed with Metafectene Pro (Biontex, Germany) according to manufacturer’s instructions.

### 2-Bromopalmitate experiments

2-Bromopalmitate (or 2-bromohexadecanoic acid, Sigma-Aldrich, Cat no. 238422) was dissolved in dimethyl sulfoxide (DMSO, Sigma-Aldrich, Cat no. D2650) to make a concentrated stock solution. Stock solution was diluted in the appropriate culture medium to a final concentration of 10µM. Cells were incubated with either 2-bromopalmitate-containing medium or control medium (the same quantity of solvent diluted in culture medium) overnight, in a humidified chamber at 37°C with 5% CO_2_. The use of ethanol as a solvent worked equally well. No particular toxicity on cultured cells was observed following incubation.

### The Acyl-Biotin Exchange (ABE) assay

Acyl-Biotin Exchange (ABE) was performed according to published protocols (Drisdel and Green, 2004; Wan et al., 2007), with slight modifications as reported in the study of Hayashi and colleagues (Hayashi et al., 2009). Briefly, the workflow of this assay involves four main steps: (i) Tissue is harvested and lysed in lysis buffer containing S-Methyl Methanethiosulfonate (MMTS, Sigma, Cat no. 64306) to block free thiol groups (-SH). (ii) Incubation with a buffer containing hydroxylamine (NH_2_OH, Fisher Scientific, Cat no. 26103) that cleaves cysteine-linked palmitate thioester bonds or with control buffer that instead of hydroxylamine contains Tris. (iii) Samples are incubated with a buffer containing biotin-HPDP (Soltec Ventures, Cat no. B106) that attaches to newly exposed cysteine thiols. Both buffers (for hydroxylamine-treated and control-treated samples) contain biotin-HPDP. Between steps i-iii, protein was precipitated by incubation with -20°C-cold 80% acetone at -20°C overnight and protein pellet was washed several times with the same solution before proceeding to next step. (iv) Biotinylated proteins are affinity-purified with streptavidin agarose resin (Thermo Scientific, Cat no. 20349). The use of neutravidin agarose resin yielded the same results. ABE samples obtained with this assay were resolved with SDS-polyacrylamide gel electrophoresis and immunoblotting with protein-specific antibodies. In this assay, the proteins detected in the plus hydroxylamine (+HA) sample but not in the minus hydroxylamine (-HA) sample (which serves as an internal negative control) represent palmitoylated proteins and collectively comprise the palmitoyl proteome. In all ABE experiments, for each tissue sample, both +HA and –HA samples were processed in parallel and the specificity of +HA signal was confirmed by the complete absence of signal in –HA sample in western blotting.

### Western blotting and quantification

Samples were mixed with Laemmli sample buffer (Laemmli, Bio-Rad, Cat no. 1610747), boiled at 95°C for 5 min, stored at -20°C and analyzed with SDS-Polyacrylamide gel electrophoresis (SDS-PAGE) and western blotting. Briefly, samples were loaded on 4-20% gradient mini-PROTEAN® TGX™ precast protein gels (Bio-Rad, 10-well Cat no. 4561094, or 15-well Cat no. 4561096) and run under standard protein electrophoresis conditions. Proteins on gels were transferred on PVDF membrane (Immobilon-P, Millipore, Cat no. IPVH00010) and blocked with 5% non-fat dry milk (Scientific, Cat no. M0841), diluted in TBS-T (TBS-Tween 20), for 1 hr at room temperature. Next, membranes were incubated with primary antibody in 1% milk/TBS-T, overnight at 4°C, with end-to-end rotation. Next day, membranes were washed with TBS-T three times and incubated with Horseradish Peroxidase (HRP)-conjugated species-specific secondary antibody in 1% milk, to a final concentration 1:10.000, for 1 hr at room temperature, with end-to-end rotation. Membranes were washed with TBS-T four times. Signal detection was performed with ECL Western Blotting Detection Reagent (GE Healthcare, Cat no. RPN2109) or ECL Prime Western Blotting Detection Reagent (GE Healthcare, Cat no. RPN2232) or Clarity Western ECL substrate (Bio-Rad, Cat no. 1705060). Different exposures were obtained for each experiment to ensure that the detected signals were in the linear range. Western blot densitometry was performed with the gel analysis tool of ImageJ. For ABE experiments, palmitoylated protein was calculated as the ratio of +HA (Hydroxylamine) signal to the respective input and subsequently all values (ratios) were expressed as a percentage of the ratio for wild-type protein (as explained in figure legends).

### Surface staining in COS-7 cells and clustering analysis

#### Surface staining

COS-7 cells were transfected with flag-tagged Nrp-2 or Nrp-1 expression plasmids. Next day, cells were incubated with anti-FLAG antibody (mouse monoclonal, Sigma, Cat no. F1804) diluted in culture medium at a final concentration 1:50, at 10°C for 20 min. Then, they were washed with fresh culture medium and fixed with ice-cold 4% paraformaldehyde in Phosphate Buffered Saline (PBS) for 10 min. Following fixation, cells were washed with PBS, blocked with 10% donkey serum and 0.1% Triton X-100 in PBS for 1 hr at room temperature and incubated with fluorescent secondary antibody (1:1000) for 1 hr at room temperature. Following antibody incubation, cells were washed with PBS and coverslips with cells were mounted on microscope slides with mounting medium (Vectashield).

#### Clustering analysis

Protein clustering on the surface of COS-7 cells was quantified with particle analysis (ImageJ). Briefly, images were converted to 8-bit, thresholded and converted to binary. Next, the image was selected and a mask of the signal was created. The “mask” image was analyzed with the “Analyze Particles” tool, with protein particles (or clusters) defined based on two parameters, size in µm^2^ (0-infinity) and circularity (0.00-1.00). Among the parameters provided in the results table for each cell analyzed are the particle count, the total area of fluorescence and the average size (area) of measured particles. Based on these, we calculate the ratio Count/Total Area, presented as “number of particles per µm^2^ of fluorescence area” in Figures 1C, 1E, 3B and S2B), which represents the degree of protein clustering; the more punctate/clustered the protein, the higher the ratio. The size of particles (Figure S2C) is an alternative measure of protein clustering; a cell with punctate protein distribution gives a low average size of protein particles, while a cell with diffuse protein distribution gives a higher average size of protein particles. Although particle analysis provides an estimate of protein distribution, it tends to underestimate differences in protein localization observed by visual inspection; thresholding softens differences between small and large particles and distinct localization patterns between Nrp-2 CS mutants are abated.

### Immunofluorescence

#### Immunofluorescence in vitro

Cells adherent on coverslips were fixed with ice-cold 4% paraformaldehyde for 10 min at room temperature, washed with PBS and blocked with 10% goat or donkey serum (depending on the primary antibody) and 0.1% Triton X-100 in PBS for 1 hr at room temperature. Next, cells were incubated with primary antibodies at 4°C overnight. Next day, they were washed with PBS and next incubated with secondary antibodies diluted 1:1000 for 1 hr at room temperature in the dark. Cells were washed four times, 10 min each, with 1x PBS and coverslips with cells were mounted on microscope slides with mounting medium (Vectashield, Vector Laboratories).

#### Immunofluorescence in vivo

Mice were anesthetized and perfused transcardially with PHEM buffer (1x PHEM: 60mM PIPES, 25 mM HEPES, 5 mM EGTA, 1 mM MgSO4; pH=6.9) followed by ice-cold PHEM buffer containing 4% paraformaldehyde and 3.7% sucrose. Brains were dissected and post-fixed with perfusion buffer (4% paraformaldehyde, 3.7% sucrose, in PHEM buffer) for 2 hrs at 4°C followed by washes with PBS and incubation with 30% sucrose in PBS at 4°C overnight. Next day, brains were embedded in NEG-50 frozen section medium following regular procedures. Sectioning was performed with a Leica cryostat at the coronal or sagittal plane at 30 µm thickness. Sections were placed on precleaned superfrost plus microscope slides (Fisher Scientific), left dry at room temperature and stored at -80°C until staining. For immunofluorescence, slides were sealed with ImmEdge hydrophobic barrier Pen (Vector Laboratories, Cat no. H-4000) and sections were blocked with 10% goat or donkey serum and 0.1% Triton X-100 in PBS for 1 hr at room temperature followed by incubation with primary antibodies diluted in 1% serum and 0.1% Triton X-100 in PBS, at 4°C overnight. Tissue was washed with PBS followed by incubation with secondary antibodies for 1 hr at room temperature. Finally, tissue was washed with PBS and covered with mounting medium (Vectashield) and a rectangular coverslip.

#### Neurofilament Staining (Floating Slice Immunofluorescence)

Mice were transcardially perfused with PBS followed by 4% paraformaldehyde in PBS. Brains were dissected and post-fixed with 4% paraformaldehyde, washed with PBS and embedded in 3% low melting point agarose. Brains were sectioned with a vibratome at the coronal plane, at 150 µm thickness, and processed for immunofluorescence floating in PBS in culture dishes. Slices were incubated with permeabilization solution (PBS, H2O, Bovine Serum Albumin, Triton X-100) at 4°C for at least 6 hrs with gentle agitation. Next, slices were incubated with anti-neurofilament (2H3) antibody (mouse monoclonal, Developmental Studies Hybridoma Bank) diluted in permeabilization solution at 4°C overnight. Next, they were washed with PBS at room temperature and subsequently incubated with secondary antibody diluted in permeabilization solution supplemented with 5% normal goat serum at 4°C overnight, washed with PBS and mounted on microscope slides with PBS. Excess PBS was removed and slices were left to dry for 30 min and subsequently covered with Fluoro Gel with DABCO (Electron Microscopy Sciences, Cat no. 17985) and a rectangular coverslip.

### Antibodies

#### Primary Antibodies (Immunofluorescence, Immunoprecipitation, Western Blotting)

Flag (mouse monoclonal, Sigma, Cat no. F1804); GFP (chicken IgY, Avés, Cat no. GFP-1020); dsRED (rabbit polyclonal, Living Colors, Clontech, Cat no. 632496); MAP2 (mouse monoclonal, Sigma, Cat no. M1406); GM-130 (rabbit monoclonal, Abcam, EP892Y, Cat no. ab52649); Neuropilin-2 (rabbit polyclonal, Cell Signaling, Cat no. 3366S); Neuropilin-2 (goat polyclonal, Research & Development, Cat no. AF2215); PlexinA3 (rabbit polyclonal, Abcam, Cat no. ab41564), Neuropilin-1 (rabbit, Abcam, Cat no. ab81321); Neuropilin-1 (goat polyclonal, R & D, Cat no. AF566); PSD-95 (mouse, NeuroMab, Cat no. 75-028); SAP102 (mouse monoclonal, NeuroMab, Cat no. 75-058); Neurofilament (2H3) (mouse monoclonal, Developmental Studies Hybridoma Bank); Ctip2 (rat monoclonal, Abcam, [25B6], Cat no. ab18465).

#### Secondary Antibodies

Immunofluorescence: CF488A donkey anti-chicken IgY (H+L) (Biotium, Cat no. 20166); Alexa Fluor 488 goat anti-mouse IgG (Life Technologies, Cat no. A11001); Alexa Fluor 488 donkey anti-goat IgG (Cat no. A11055); Alexa Fluor 555 donkey anti-rabbit IgG (Life Technologies, Cat no. A-31572); Alexa Fluor 546 donkey anti-mouse IgG (ThermoFisher Scientific, Cat no. A10036); Alexa Fluor 555 goat anti-rabbit IgG (Cat no. A-21428); DyLight 649 donkey anti-mouse IgG (Jackson ImmunoResearch); Alexa Fluor 405 goat anti-rabbit IgG (ThermoFisher Scientific, Cat no. A-31556); Alexa Fluor 647 goat anti-chicken IgG (ThermoFisher Scietific, Cat no. A-21449); Alexa Fluor 647 donkey anti-rabbit IgG (ThermoFisher Scientific, Cat no. A-31573); Alexa Fluor 647 donkey anti-rat IgG (Abcam, Cat no. ab150155).

Western Blotting: Horseradish peroxidase (HRP)-conjugated antibody directed against the host species of the primary antibody.

### Nrp-2–Golgi association analysis

#### Immunofluorescence

E14.5 *Nrp-2^-/-^* cortical neurons in culture were transfected with pHluorin-tagged Nrp-2 expression plasmids and at DIV17 they were stained with a GFP antibody to detect Nrp-2 (total, surface and intracellular) and a GM130 antibody to visualize cis-Golgi. Neurons were imaged with an upright LSM 700 confocal microscope (Zeiss) with the acquisition of single images.

#### Quantitative Assessment of Colocalization

Performed with ImageJ. The Nrp-2 and Golgi (GM130) channels were split and thresholded creating binary images (a different threshold was used for each of the two channels, while thresholds were kept constant throughout the analysis). The area of Nrp-2 signal (A_Nrp-2_) and Golgi signal (A_Golgi_) was selected and measured (µm^2^). The colocalization plugin was used to give the colocalized image that was used to create a mask that represents the colocalized area (A_Nrp-2/Golgi_), which was then selected and measured. The ratio of the colocalized area to the Nrp-2 area (A _Nrp-2/Golgi_/A_Nrp-2_) represents the fraction of Nrp-2 that is associated with Golgi. This ratio was further normalized to the total Golgi area (A_Golgi_) to take into account differences in the abundance of Golgi membranes between neurons.

#### Golgi Isolation

Performed on wild-type mouse whole brain according to *Current Protocols in Cell Biology* (Unit 3.9, Basic Protocol 2). Mouse brain was harvested and gently homogenized with a Potter-Elvehjem pestle to avoid Golgi stack fragmentation. Golgi stacks were isolated by flotation through a discontinuous sucrose gradient. Briefly, light mitochondrial pellet was resuspended in 2 ml buffer and mixed with 8 ml 2.0 M sucrose so that final sucrose concentration is about 1.55 M. 5 ml of this mix was transferred to a 17-ml ultracentrifuge tube (Beckman) and overlayed with the following sucrose solutions: 4 ml 1.33 M sucrose, 2 ml 1.2 M sucrose, 2 ml 1.1 M sucrose, 2 ml 0.77 M sucrose, 0.25 M sucrose to fill the tube. Gradients were centrifuged at 100,000 x g at 4°C for 1 hr. The following fractions were collected: 1.55 M, 1.55 M/1.33 M, 1.33 M, 1.1 M/1.2 M, intervening fraction, 0.77 M/1.1 M. Samples were prepared and stored for SDS-PAGE and immunoblotting.

### Live Imaging

Live imaging was performed to visualize cell surface localization of pHluorin-tagged Nrp-2 in primary cortical neurons or Neuro2A cells. The pHluorin epitope tag serves as a robust fluorescent reporter of protein expressed on the cell surface because of its sensitivity to pH. Cells were cultured in individual glass bottom dishes (MatTek Life Sciences, Cat no. P35G-0-14-C) precoated with poly-D-lysine and transfected with appropriate plasmids. One or two days after transfection, they were imaged live at an inverted LSM 700 confocal microscope (Zeiss) with a 40X oil lens.

### Fluorescence Recovery After Photobleaching (FRAP)

Wild-type cortical neurons were cultured and plated on glass bottom dishes precoated with poly-D-lysine. Between DIV5 and DIV8 neurons were transfected with the indicated pHluorin-Nrp-2-IRES-dsRED expression plasmids with the method of Lipofectamine. A few days later, they were imaged with an inverted LSM 700 microscope (Zeiss) with an incubation chamber (PeCon) (temperature: 37°C, CO_2_: 5%), using a 63X NA1.4 oil immersion lens. For the performance of fluorescence recovery after photobleaching (FRAP), we used the following protocol: a region of interest (ROI) containing the main dendritic process was selected for FRAP analysis. Images were acquired every 30 s; 5 images were acquired prior to bleaching (prebleach time: 2.5 min) and 31 additional images were acquired post-bleaching (total post-bleach time: 15.5 min). Analysis: Fluorescence intensity of time-lapse images was determined by measuring the fluorescence intensity within the ROI and correcting for fluorescence decay and for background fluorescence.

### Immunoprecipitation in heterologous cells

Cell growth medium was aspirated and lysis buffer (TNE buffer: NP-40, Tris, EDTA, NaCl, protease inhibitor cocktail) was added in cells in the culture plate. Cells were scraped off, mechanically lysed on ice by passage through 1 ml syringes and incubated with rotation at 4°C for 1 hr. Samples were precleared with Protein A/G agarose resin (Pierce, ThermoFisher Scientific, Cat no. 20421) at 4°C for 2 hrs. Next, after the resin was discarded, a small fraction of each sample was kept as input and the remaining samples were incubated with the appropriate antibody at 4°C overnight. Next, samples were incubated with protein A/G agarose with end-over-end rotation at 4°C for 2 hrs. Resin was washed twice with high-salt buffer (500 mM NaCl) and twice with low-salt buffer (150 mM NaCl). Proteins were eluted from resin with Laemmli sample buffer diluted in TNE buffer, boiled at 95°C for 5 min and analyzed with SDS-PAGE and western blotting.

### Alkaline Phosphatase (AP)−fused ligand production

Alkaline Phosphatase (AP)-tagged Sema3A and Sema3F or AP expression plasmids were transfected into 293T cells with the method of Lipofectamine. Cells were allowed to secrete the ligand for 2-5 days. The culture supernatant was collected and transferred to a 50 ml MW filter tube (Centricon filters, Millipore) and was centrifuged at 35 rpm for 10 min at 4°C. A 50K filter was used for AP concentration and a 100K filter was used for concentration of Sema3A-AP or Sema3F-AP. The liquid above the filter was transferred to a new tube and ligand concentration was determined with a spectrophotometer by measuring AP activity by the change in absorbance at OD 405 nm. This figure was then converted to nM. The concentrated ligand was aliquoted and stored at -80°C. For each experiment, a new aliquot was thawed and used for culture treatment to avoid protein decay from thawing and refreezing.

### *Nrp-2^-/-^* rescue experiments

For in vitro rescue, E14.5 *Nrp-2^-/-^* primary cortical neurons were transfected at DIV7 with the indicated plasmids. At DIV21 neurons were treated with 5 nM Sema3F-AP or AP for 6 hrs and afterwards they were processed for immunofluorescence. Confocal images (stacks) were taken with an LSM 700 with a 63X oil lens. Analysis was performed with ImageJ and dendritic spines were counted along the 0-50 µm apical dendritic segment from the cell body. For in vivo rescue, *in utero* electroporation was performed on E13.5 *Nrp-2^F/-^* embryos of timed-pregnant females (detailed below). *Nrp-2^-/-^* mice were crossed with *Nrp-2^F/F^*mice so that all embryos are *Nrp-2^F/-^*.

### Neuron labeling in vivo

#### In Utero Electroporation

Performed on E13.5 *Nrp-2^F/-^* embryos of timed-pregnant females according to the published protocol (Saito, 2006). Briefly, anesthesia was delivered with intraperitoneal injection of sodium pentobarbital and, upon anesthetization, a small vertical incision was made on the abdominal cavity and embryos were gently pulled out. DNA was microinjected in the lateral ventricle with a fine and polished glass capillary tube placed in a mouth-controlled pipette, followed by the administration of 5 electric pulses of 30-35 V and 50 ms each with inter-pulse interval 950 ms, by the use of forceps-type electrodes. Embryos were placed back in the mother’s abdomen, the incision was closed with silk sutures or surgical staples and the mouse was placed back in the cage on a slide warmer, with hydrogel next to it, and monitored until full recovery. After the surgery and until pup delivery the pregnant female was kept in the High Risk Mouse Room. Newborn mice were sacrificed on P28 and brains were examined for direct fluorescence with a stereoscope. Electroporated brains were processed for immunofluorescence that included triple staining for EGFP, dsRED and Ctip2; the latter to confirm that labeled neurons used for phenotypic analysis are Ctip2^+^ (layer V pyramidal neurons).

#### Genetic Labeling

The Thy1-EGFP-m line was crossed to *DHHC15^-/-^* mice for labeling of layer V cortical neurons. *DHHC15^+/-^*;*Thy1-EGFP* mice were crossed to generate litter- and age-matched wild-type and *DHHC15^-/-^* mice expressing Thy1-EGFP. Thy1-EGFP^+^ brains were subjected to double staining against EGFP (to visualize neuronal architecture) and Ctip2 (to confirm that EGFP-labeled neurons used for phenotypic analysis are Ctip2^+^).

#### Golgi Staining

Performed with the use of Rapid GolgiStain kit (FD Neurotechnologies, Cat no. PK401) according to manufacturer’s protocol. Briefly, brains were dissected and incubated with impregnation solution A+B for 12 days in darkness at room temperature (solution was replaced by fresh the second day). After 12 days, brains were incubated with solution C for 4 days in darkness at 4°C (solution C was replaced by fresh the second day). Brains were embedded in NEG-50 frozen section medium following regular procedures and sectioned with a cryostat (Leica) at the sagittal plane at 100 µm thickness. Sections were transferred on gelatin-coated superfrost plus slides, left to dry in the dark overnight and next day they were stained according to the protocol. Neurons were imaged with acquisition of stacks using a DIC at an inverted LSM 700 confocal microscope (Zeiss), with a 20X lens for visualization of dendrites and a 63X oil lens for visualization of dendritic spines.

### Phenotypic analysis

#### Dendritic spine assay and analysis

For dendritic spine assessment in vitro, neurons were cultured on glass coverslips until DIV21 to allow spine formation. Between DIV7 and DIV9 they were transfected with pCIG2-ires-EGFP–expressing plasmids and at DIV21 they were treated with 5 nM Sema3F-AP or 5 nM AP (control) in neuron growth medium for 6 hrs in the tissue culture incubator (37°C, 5% CO_2_). Subsequently, neurons were fixed with ice-cold 4% paraformaldehyde for 10 min at room temperature and subsequently subjected to EGFP immunofluorescence. For both in vitro and in vivo analysis, EGFP-expressing neurons were imaged with an upright laser scanning confocal microscope (LSM 700, Zeiss) with a 63X oil lens by the acquisition of Z-stacks. For spine analysis, the 3D projection was calculated and spines were counted along the proximal 50 µm of the apical dendrite (relative to the cell body). All subtypes of dendritic spines (stubby, mushroom, filopodia-like) were counted.

#### Dendritic arborization assay and analysis

For dendritic arborization assessment in vitro, neurons were cultured on glass coverslips until DIV12 to allow dendritic elaboration. At DIV12 primary cortical neurons were treated with 5 nM Sema3A-AP or 5 nM AP (control) for 6 hrs in neuron growth medium in the tissue culture incubator (37°C, 5% CO_2_). After treatment, they were fixed with 4% paraformaldehyde and subjected to immunofluorescence with an antibody directed against the somatodendritic marker MAP2. For both in vitro and in vivo analysis, imaging was performed at an LSM 700 upright confocal microscope (Zeiss) with a 40X oil lens with the acquisition of Z-stacks. The calculated 3D projection was thresholded and any background fluorescence or neighboring neurons interfering with the analysis were erased with the paintbrush tool. The point-selection was placed at the center of the cell body and Sholl analysis (ImageJ) was run to count the dendritic processes at various distances from the cell body, ranging from 0 µm to 100 µm or 150 µm. For phenotypic analysis, efforts were made so that analyzed mice of different genotypes are sex-matched so that any sex-related variability is not a confounding factor for the interpretation of experimental data.

### Statistical analysis and softwares

Statistical significance and the size of samples analyzed are presented in Figures and Figure Legends. Image analysis was performed using ImageJ software with appropriate plugins (Rasband, W.S., ImageJ, U.S. National Institutes of Health, Bethesda, Maryland, USA, https://imagej.nih.gov/ij/, 1997-2016). The prediction of neuropilin palmitoylation sites was performed with the CSS-Palm 3.0 software, which is freely available (Ren et al., 2008). The ClustalW software used for conservation analysis of Nrp-2 and Nrp-1 amino acid sequences is freely available. Details for the analysis of individual biochemical and phenotypic experiments are provided in separate sections. Data were collected in Excel and GraphPad Prism. Generation of all graphs and statistical analyses were performed in GraphPad Prism. The GraphPad Prism software was purchased. Figure assembly and construction of Figures 7E and 7F were carried out in Adobe Illustrator. Statistical analysis tests include two-tailed t test and Analysis of Variance (ANOVA) followed by the appropriate post-hoc test for multiple comparisons (as mentioned in figure legends). Statistical significance is defined as: *p < 0.05; **p < 0.01, ***p < 0.001, ****p < 0.0001, ns: not significant. Exact p values are presented either on graphs or in figure legends.

### Data availability

All quantifications and the western blots reported in the manuscript are included in the manuscript and in the supporting files uploaded during submission. During review, additional raw data, including image files and western blots used for quantifications, can be provided upon request. Upon manuscript acceptance, a repository with all raw data and source files will be generated and will become accessible to all scientists via an accession number.

**Figure 2—figure supplement 1.**
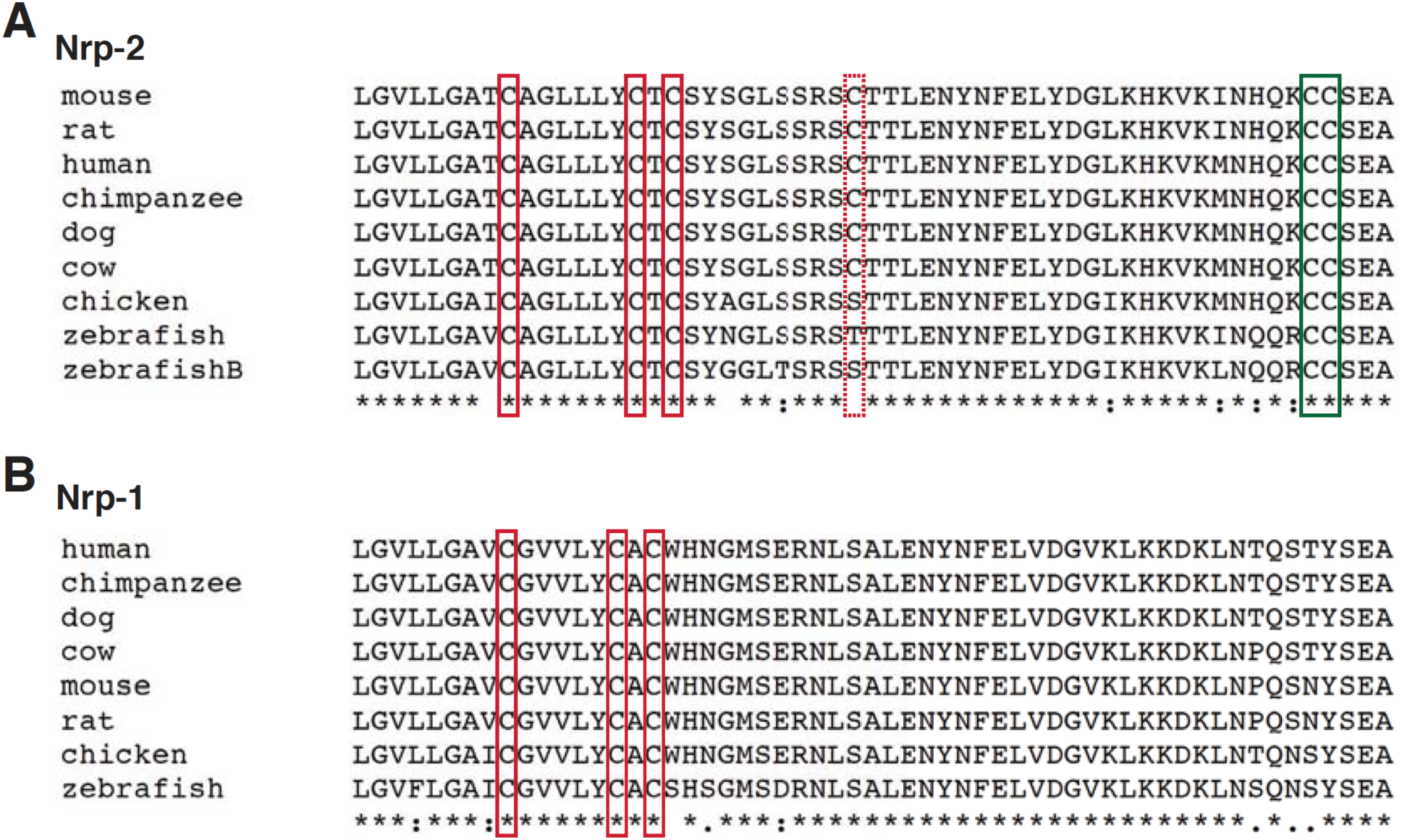
Conserved cysteine residues lie in the transmembrane and cytoplasmic domains of neuropilins. **(A)** Nrp-2 cross-species amino acid sequence alignment of transmembrane and cytoplasmic domains, by the use of ClustalW software, reveals highly conserved cysteine residues across species. The three transmembrane/juxtamembrane cysteines (surrounded by red boxes) and the two C-terminal cysteines (surrounded by green box) are highly conserved, while the cytoplasmic cysteine in the middle (surrounded by dashed red box) is partially conserved. **(B)** Nrp-1 cross-species amino acid sequence alignment of transmembrane and cytoplasmic domains, as mentioned above, reveals highly conserved cysteine residues (surrounded by red boxes) across species in its transmembrane/juxtamembrane domains.

**Figure 3—figure supplement 1.**
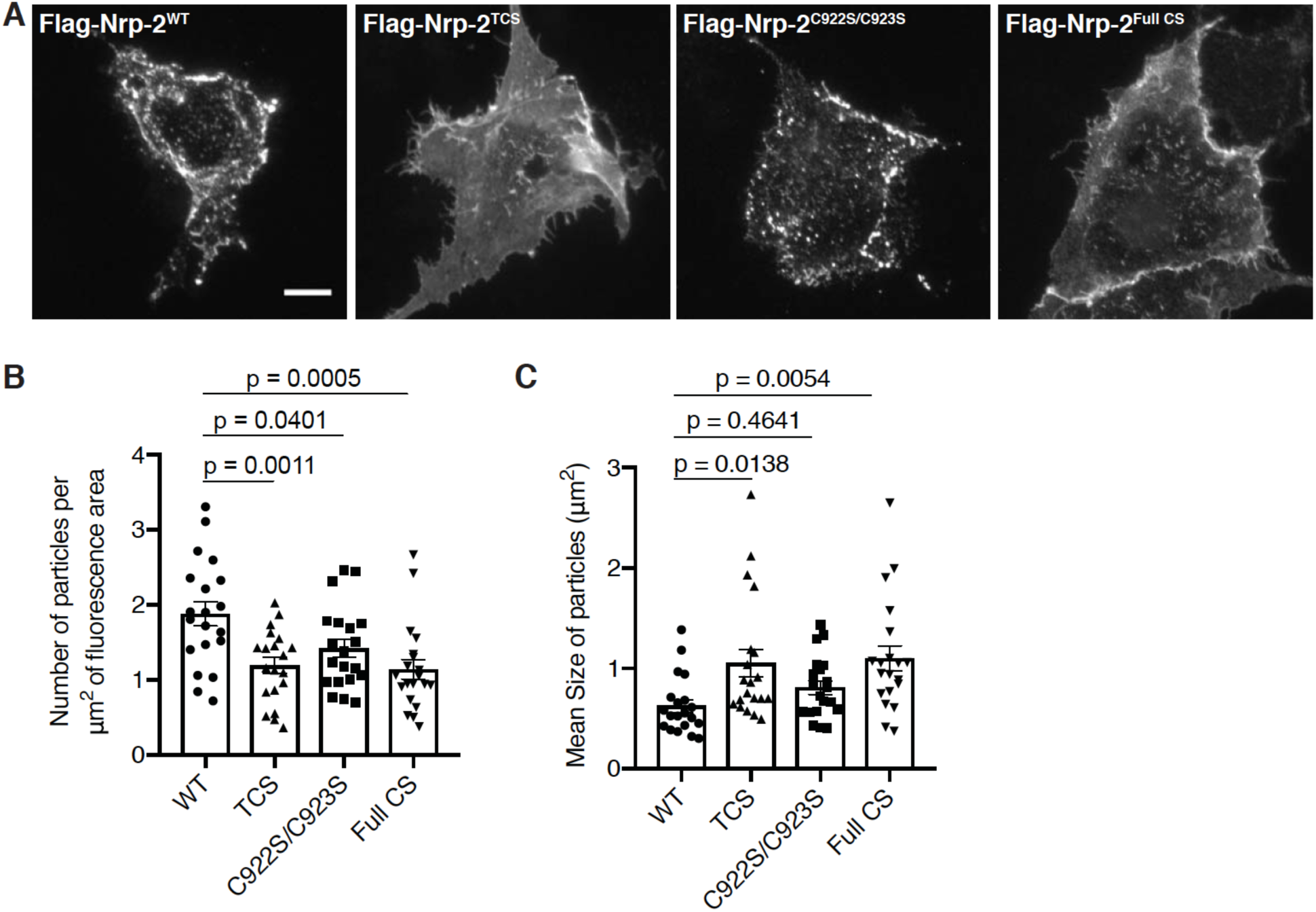
Distinct requirements for Nrp-2 palmitoyl acceptor cysteines in Nrp-2 cell surface distribution in COS-7 cells. **(A-C)** Cell surface localization of Nrp-2 protein in heterologous cells. (A) Panels show COS-7 cells expressing flag-tagged Nrp-2 wild-type (WT) or various CS mutants, subjected to Nrp-2 surface staining with a flag antibody. Scale bar, 15 µm. (**B, C**) Quantification of protein clustering with particle analysis (see Materials and Methods), presented as number of particles (clusters) per µm^2^ of fluorescence area (B) and as particle size (area in µm^2^); each cell gave an average particle size (**C**). Data are plotted in scatter dot plots with mean ± SEM. WT and C922S/C923S Nrp-2 are distributed in the form of smaller clusters (puncta), whereas TCS and Full CS localize on the surface as larger clusters indicative of a diffuse distribution pattern. One-way ANOVA followed by Dunnett’s test for multiple comparisons; n = 20 cells analyzed for each plasmid.

**Figure 3—figure supplement 2.**
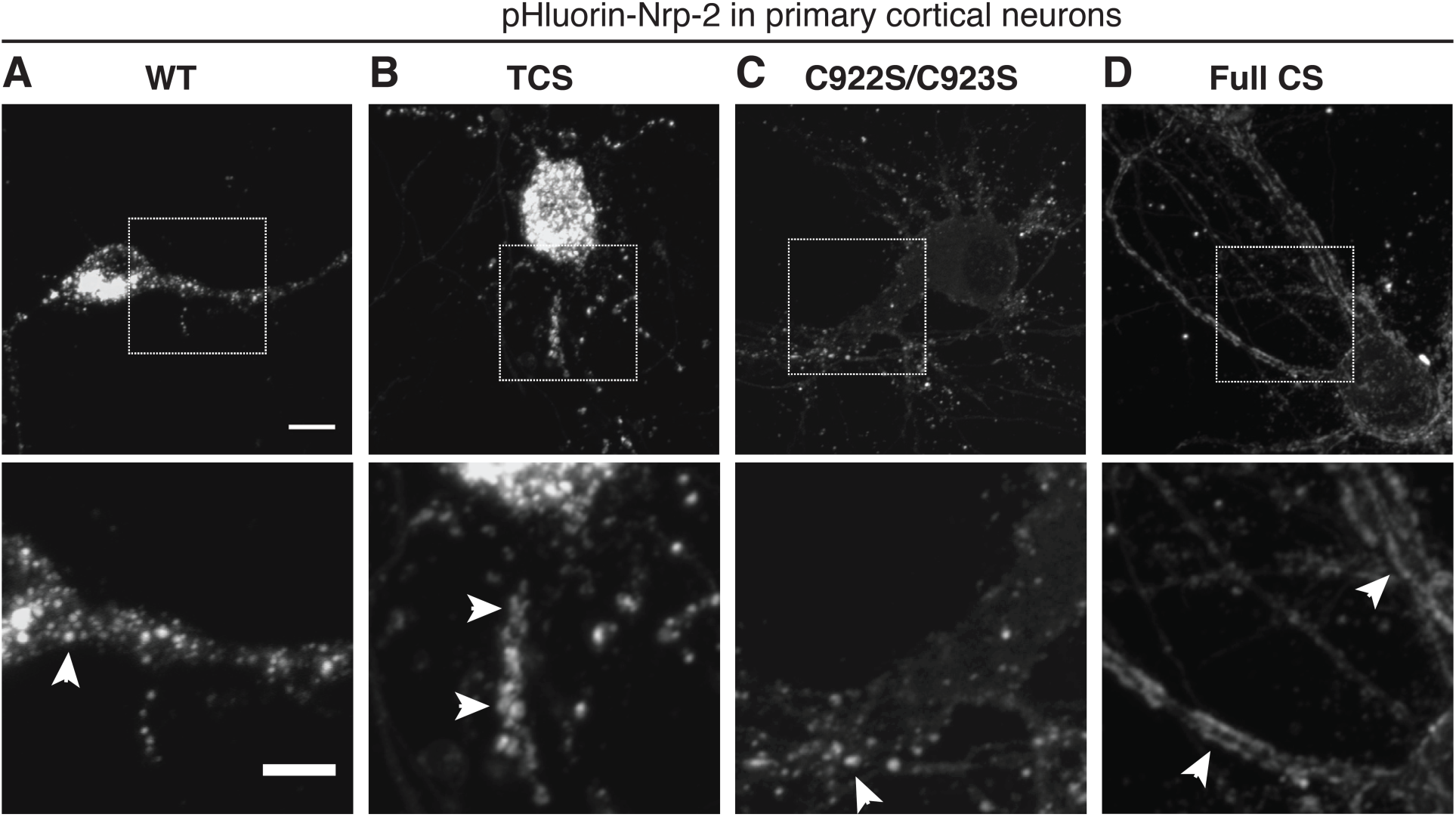
Severe defects in the cell surface localization of palmitoylation-deficient Nrp-2 in primary cortical neurons. **(A-D)** Surface localization of Nrp-2 lacking palmitoyl acceptor cysteines in cortical neuron cultures. Primary cortical neurons were transfected with pHluorin-tagged Nrp-2 WT or CS mutants and at DIV16 they were imaged live to visualize protein localized on the plasma membrane. Top panels show representative images of neurons expressing each plasmid; bottom panels show the zoomed-in view of the areas surrounded by rectangles in top panels. WT (A) and C-terminal CS (C) Nrp-2 proteins are punctate (arrowheads mark puncta), whereas Nrp-2 TCS (B) is mostly distributed in the form of larger clusters resembling clumps (arrowheads) and to a much lesser extent in the form of puncta. Outstandingly, palmitoylation-deficient Nrp-2, Full CS (D), has almost completely lost clustering and diffuses over the entire cell surface, making a pattern reminiscent of “tram track” (marked by arrowheads). Scale bars: top panels, 8 µm; bottom panels, 5 µm.

**Figure 3—figure supplement 3.**
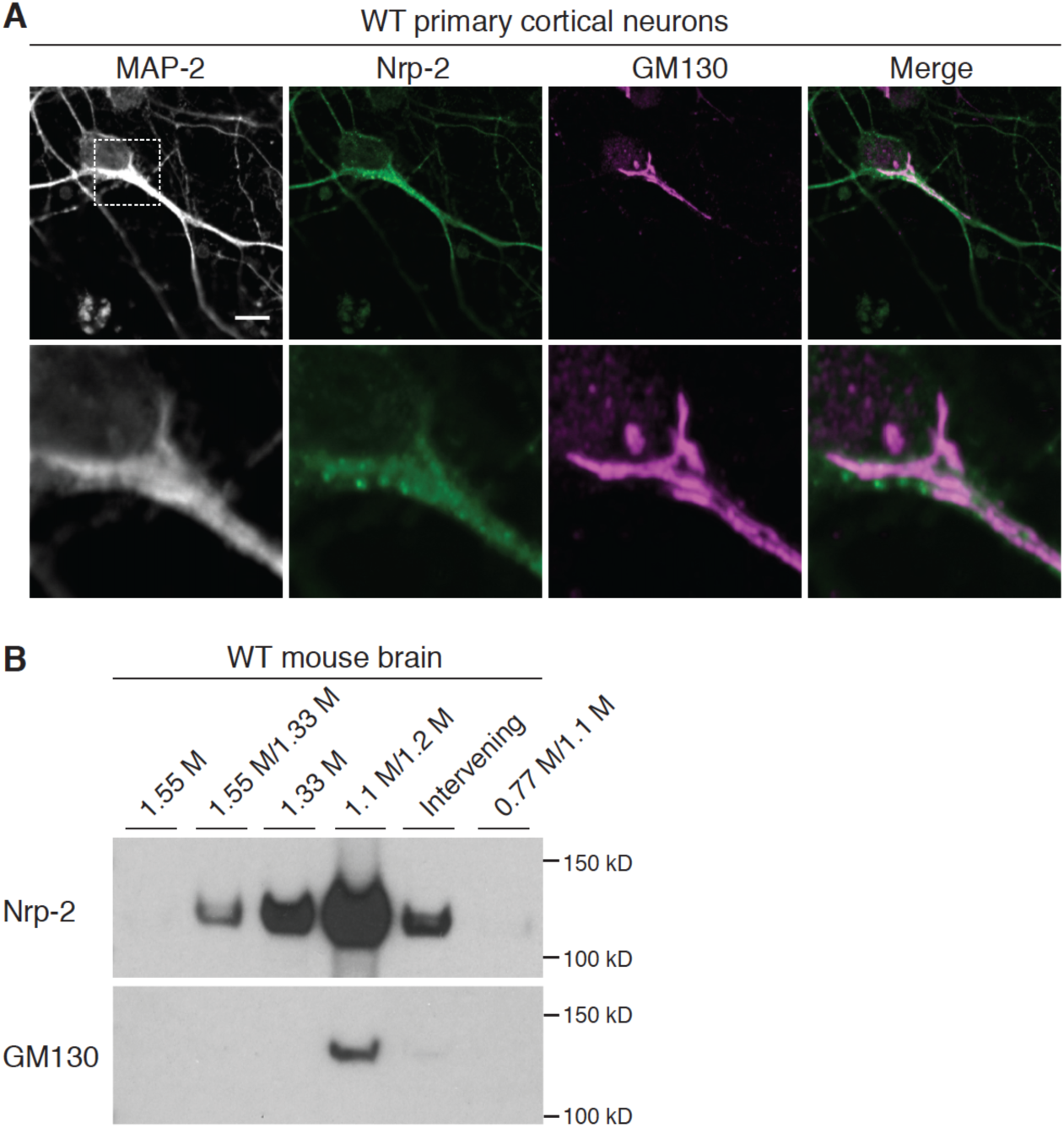
Nrp-2 is enriched in the Golgi apparatus in neural tissue. **(A)** Wild-type primary cortical neurons immunostained with antibodies against Nrp-2 and the *cis*-Golgi marker GM130. Endogenous Nrp-2 robustly colocalizes with somatic Golgi and dendritic Golgi membranes, as shown in the overlaid image (merge). Bottom row shows a zoomed-in view of the area surrounded by the dashed rectangle. Scale bar, 7 µm. **(B)** Golgi isolation preparations made from adult mouse whole brain, analyzed with western blotting using antibodies directed against Nrp-2 and GM130. Endogenous Nrp-2 is highly enriched in the GM130-positive fraction (1.1M/1.2M).

**Figure 3—figure supplement 4.**
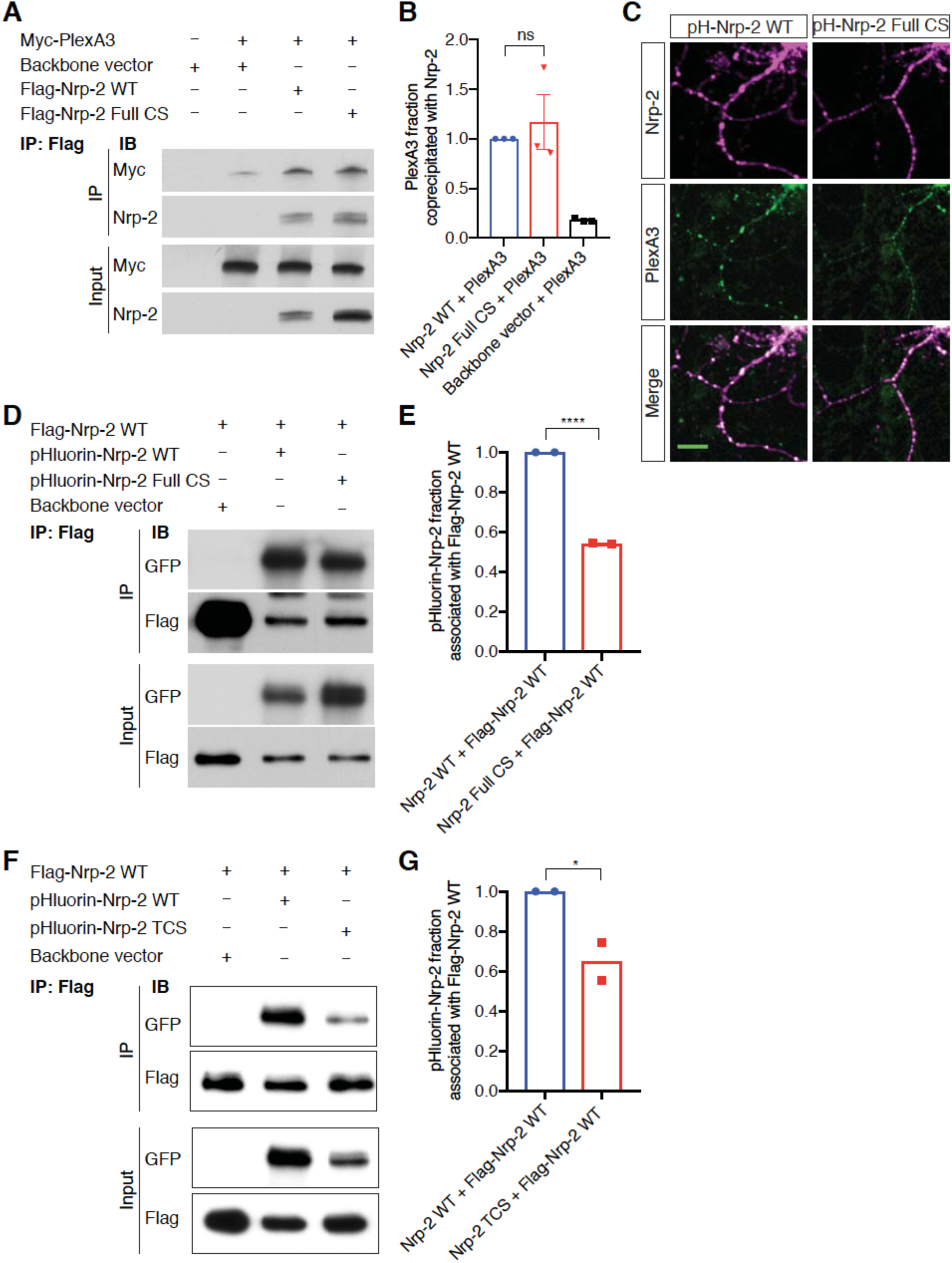
Nrp-2 palmitoyl acceptor cysteines are not required for Nrp-2/PlexA3 association but are required for proper Nrp-2 homooligomerization. (**A, B**) Effects of Nrp-2 palmitoyl acceptor cysteines on Nrp-2/PlexA3 complex formation, assessed by co-immunoprecipitation experiments. 293T cells were transfected with myc-tagged PlexA3 and either pHluorin-tagged WT Nrp-2 or pHluorin-tagged Full CS Nrp-2 or backbone vector. Nrp-2 immunoprecipitation was performed with a GFP antibody. (A) Immunoblotting against the myc epitope tag shows that PlexA3 is co-immunoprecipitated with wild-type or Full CS Nrp-2 to similar degrees. (B) Quantification of PlexA3 fraction immunoprecipitated, calculated as the ratio IP/Input. This fraction for WT Nrp-2 samples is set at 100% and the fraction for Full CS Nrp-2 samples is expressed relative to that. Data are plotted in dot plots with mean ± SEM. Two-tailed t test, p = 0.5781; ns, not significant; n = 3 experiments. (**C**) Association of PlexA3 with Nrp-2, in DIV21 wild-type primary cortical neurons transfected with pHluorin-tagged Nrp-2 WT or Full CS and subjected to immunofluorescence against GFP (Nrp-2 epitope tag) and endogenous PlexA3. Both WT and palmitoylation-deficient Nrp-2 proteins (magenta) display strong association with PlexA3 (green), shown as white in overlaid images (merge). Scale bar, 20 µm. (**D, E**) Assessment of cysteines’ effects in the ability of Nrp-2 to homomerize. (D) Western blots of co-immunoprecipitation experiments in 293T cells co-transfected with flag-tagged WT Nrp-2 and either pHluorin-tagged WT Nrp-2 or pHluorin-tagged Full CS Nrp-2 or backbone vector. Flag-tagged WT Nrp-2 was immunoprecipitated with a flag antibody and samples were immunoblotted with flag or GFP antibody. (**E**) Quantification of the fraction of pHluorin-tagged Nrp-2 (WT or Full CS) that is associated with flag-tagged Nrp-2 WT. This fraction is set at 1.0 for WT and Full CS is expressed as a percentage of that. Full CS Nrp-2 associates with WT Nrp-2 to a significantly lesser extent than WT Nrp-2 does. Data are plotted in dot plots with mean ± SEM, Two-tailed t test, ****p < 0.0001, n = 2 experiments. (**F, G**) Assessment of cysteines’ effects in the ability of Nrp-2 to homomerize. (F) Western blots of co-immunoprecipitation experiments in 293T cells co-transfected with flag-tagged WT Nrp-2 and either pHluorin-tagged WT Nrp-2 or pHluorin-tagged TCS Nrp-2 or backbone vector. Flag-tagged WT Nrp-2 was immunoprecipitated with a flag antibody and samples were immunoblotted with flag or GFP antibody. (**G**) Quantification of the fraction of pHluorin-tagged Nrp-2 (WT or TCS) that is associated with flag-tagged Nrp-2 WT. This fraction is set at 1.0 for WT and TCS is expressed as a percentage of that. Data are plotted in dot plots with mean ± SEM. TCS Nrp-2 associates with WT Nrp-2 less than WT Nrp-2 does. One-tailed t test, *p = 0.0317, n = 2 experiments.

**Figure 4—figure supplement 1.**
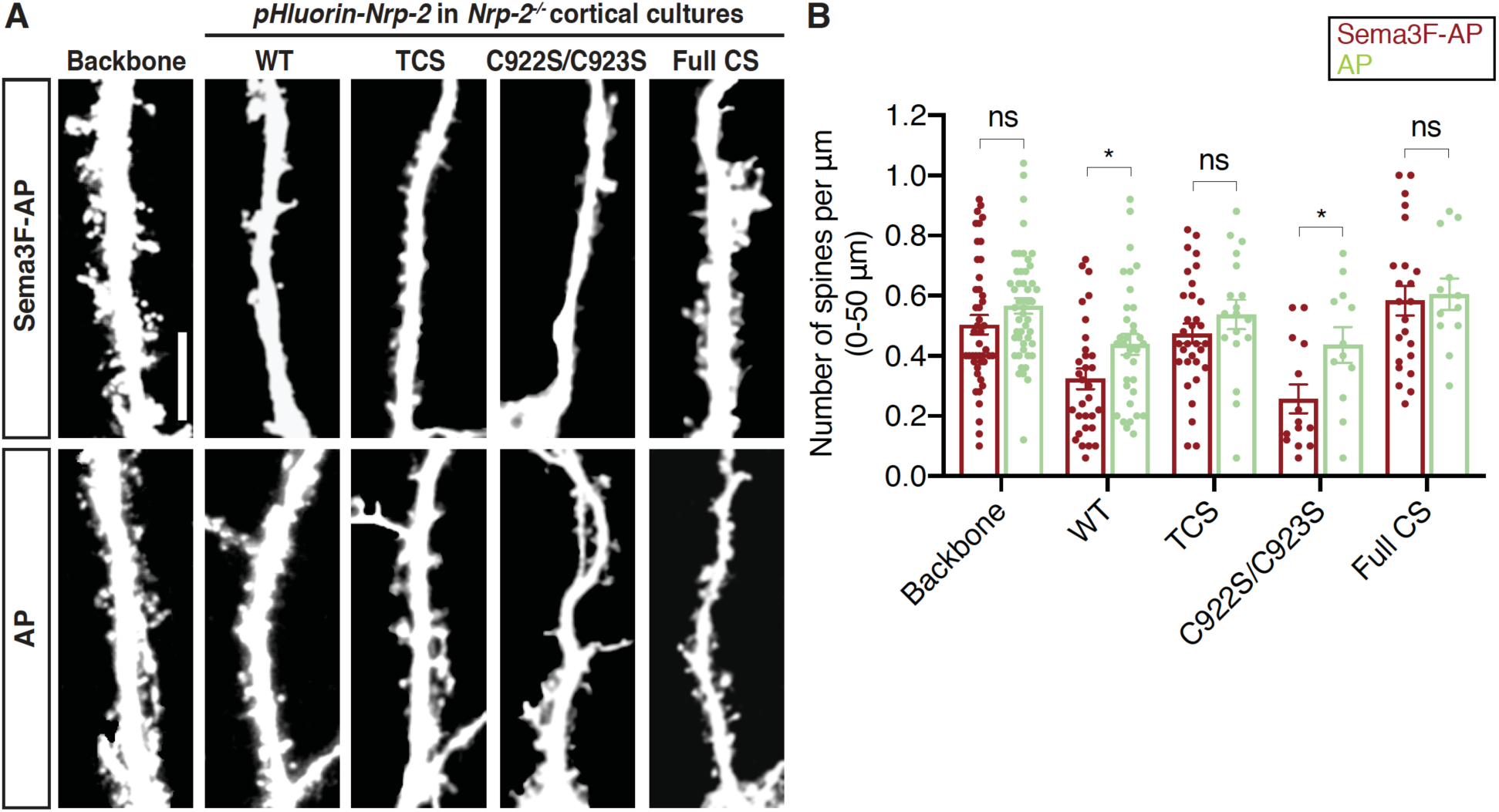
Sema3F/Nrp-2-dependent dendritic spine pruning in deep layer cortical neurons depends on distinct Nrp-2 cysteine clusters. (**A, B**) Rescuing in vitro the *Nrp-2^-/-^* dendritic spine phenotype to assess the role of different Nrp-2 cysteine motifs in Sema3F/Nrp-2-dependent dendritic spine retraction. E14.5 *Nrp-2^-/-^* primary cortical neurons were transfected with various pHluorin-tagged *Nrp-2-ires-dsRED* expression plasmids, including WT, TCS, C922S/C923S and Full CS. At DIV21 they were treated with 5 nM Sema3F-AP or AP, for 6 hrs, and next subjected to immunofluorescence. (A) Representative images of neurons (3D projection of confocal stack) expressing the indicated plasmid treated with either Sema3F-AP or AP. Scale bar, 10 µm. (B) Quantification of dendritic spines, counted along the 0-50 µm apical dendritic segment and presented as number of spines per µm. Data are plotted in scatter dot plots with mean ± SEM. Graph shows two columns for each plasmid, one for spine density in AP-treated neurons and one for spine density in Sema3F-AP-treated neurons. The rescue ability of each Nrp-2 protein is evidenced by the ability of *Nrp-2^-/-^* neurons expressing each Nrp-2 plasmid to cause spine pruning in response to Sema3F-AP as compared to AP treatment (control). Sema3F-AP treated: Backbone, n = 45; WT, n = 31; TCS, n = 32; C922S/C923S, n = 14; Full CS, n = 23. AP treated: Backbone, n = 48; WT, n = 34; TCS, n = 18; C922S/C923S, n = 12; Full CS, n = 12, where n is the number of neurons analyzed for each condition (Two-tailed t test; Sema3F-AP vs AP for each tested plasmid: Backbone, p = 0.1321; WT, p = 0.0222; TCS, p = 0.2729; C992S/C923S; p = 0.0251; Full CS, p = 0.7842; ns, not significant, p ≥ 0.05; ANOVA could not be reliably performed because of the highly unequal sizes (number of neurons) of different conditions).

**Figure 6—figure supplement 1.**
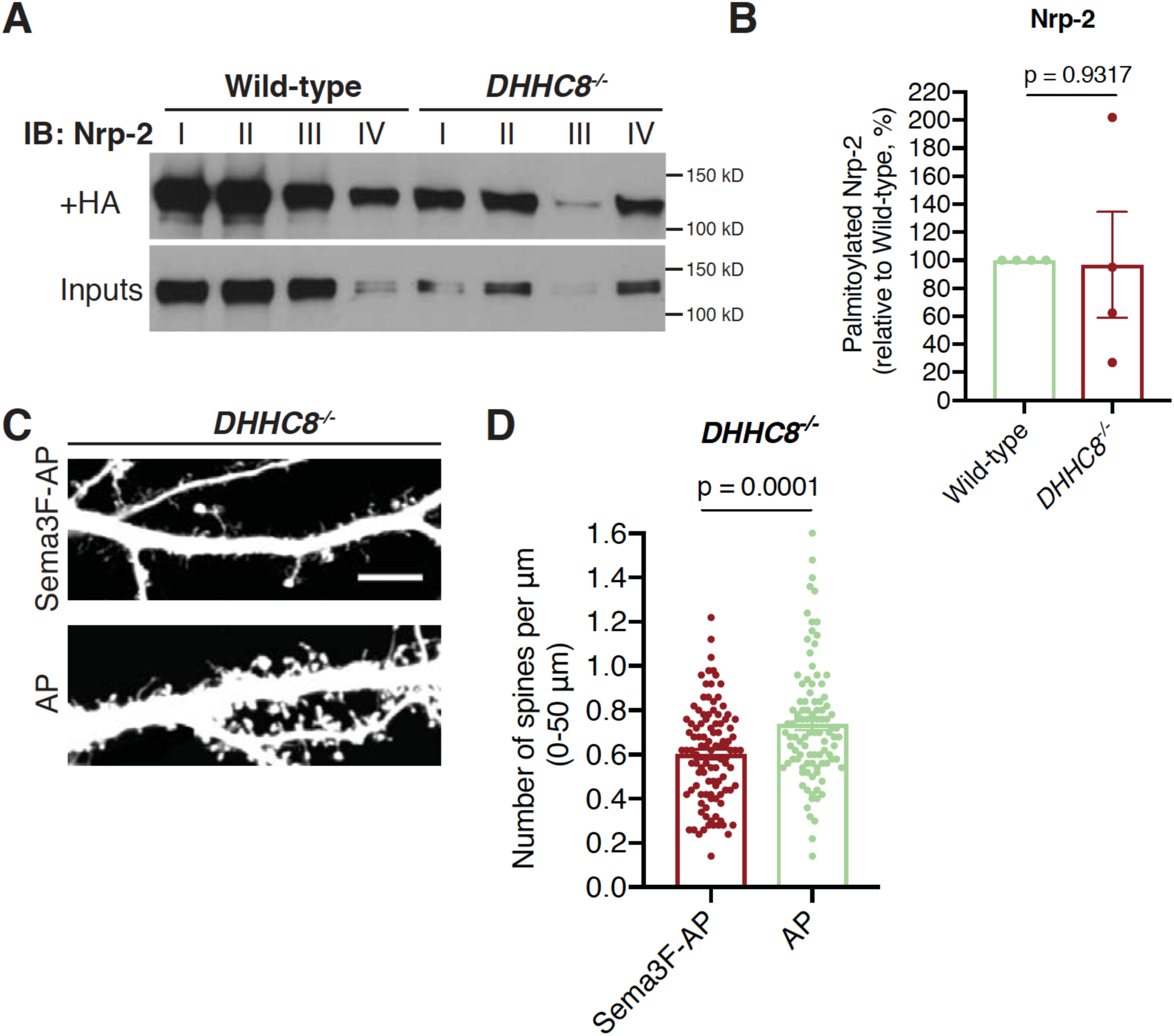
Nrp-2 is not a substrate for the palmitoyl acyltransferase DHHC8 in deep layer primary cortical neurons. **(A, B)** ABE performed on E14.5 DIV12 wild-type C57BL/6 or *DHHC8^-/-^* primary cortical neurons. (A) Nrp-2 immunoblotting of palmitoylated (+HA) and input samples; I, II, III, IV represent biological replicates. (B) Quantification of palmitoylated Nrp-2 levels, calculated as the ratio of +HA to the respective input; this ratio for *DHHC15*^-/-^ neurons is expressed as a percentage of that in wild-type (set at 100%). Pooled data from all four independent experiments are plotted in scatter dot plot with mean ± SEM. These experiments reveal no consistent difference in Nrp-2 palmitoylation between wild-type and *DHHC8^-/-^* cultured neurons. Two-tailed t test. **(C, D)** Sema3F causes dendritic spine pruning on DIV21 *DHHC8^-/-^* deep layer cortical neurons in culture. (C) Representative images are shown for each treatment group (Sema3F-AP and AP control group) and constitute the 3D projection of confocal stacks. Scale bar, 7 µm. (D) Quantification of spines along the proximal 50 µm on the largest dendrite (extending distally from the cell body), presented as number of spines per µm and plotted in scatter dot plot with mean ± SEM (2 independent experiments). Two-tailed t test; Sema3F-AP: n = 103, AP: n = 99, where n is the number of neurons analyzed.

**Figure 7—figure supplement 1.**
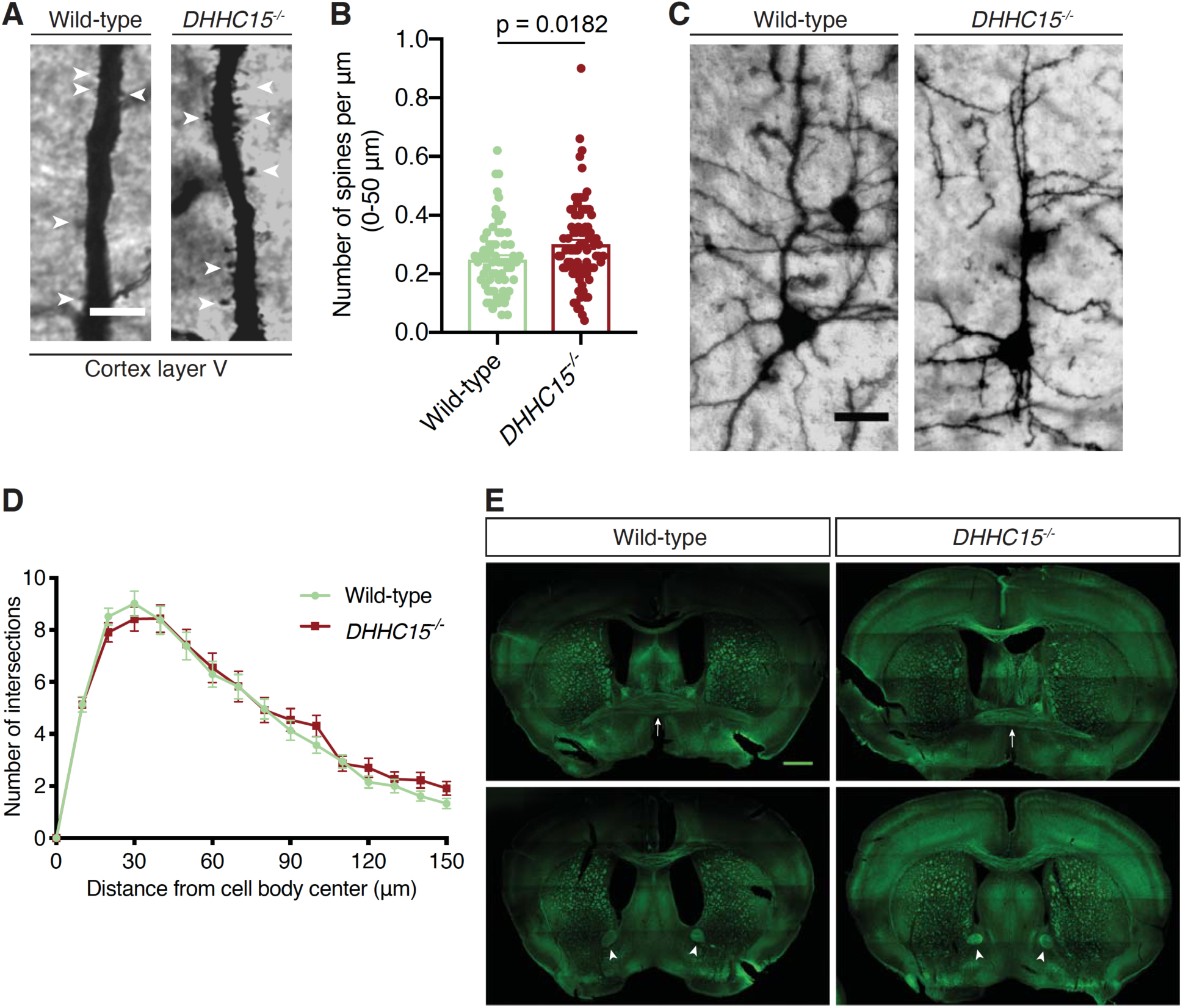
Selective effects of DHHC15 on Nrp-2-dependent developmental processes in the mouse brain. **(A, B)** Analysis of dendritic spine density on layer V cortical pyramidal neurons of wild-type and *DHHC15^-/-^* brains labeled with Golgi staining. (A) Representative images are shown for each genotype and constitute the 3D projection of confocal stacks. White arrowheads point to dendritic spines. Scale bar, 7 µm. (B) Quantification of dendritic spines, counted along the proximal 50 µm (relative to the cell body) on neurons’ apical dendrite. Data from independent experiments (3 pairs of mice-littermates, Wild-type vs *DHHC15^-/-^*) are plotted in scatter dot plot with mean ± SEM. Two-tailed t test; Wild-type, n = 65; *DHHC15^-/-^*, n = 85; where n is the number of neurons analyzed. **(C, D)** Assessment of dendritic arborization of layer V cortical pyramidal neurons of wild-type and *DHHC15^-/-^* brains labeled with Golgi staining. (C) Representative images are shown for each genotype and represent the 3D projection of confocal stacks. Scale bar, 30 µm. (D) Quantification of dendritic arborization with Sholl analysis reveals no significant difference in dendritic arbor complexity between wild-type and *DHHC15^-/-^* layer V pyramidal neurons of the cerebral cortex. Data from independent experiments (3 pairs of mice-littermates, Wild-type vs *DHHC15^-/-^*) are plotted as mean ± SEM. Multiple t tests; Wild-type, n = 72; *DHHC15^-/-^*, n = 64, where n is the number of neurons analyzed. **(E)** Neurofilament (2H3) staining on coronal brain sections of adult wild-type C57BL/6 and *DHHC15* homozygous mutant mice reveals intact midline crossing fibers (marked by white arrows) and anterior limb (marked by white arrowheads) of the anterior commissure in both wild-type and mutant mice (observed in all mice analyzed, n = 3 mice per genotype). Images are taken with tile scan (horizontal and vertical lines represent artifacts of tile scan). Scale bar, 800 µm.

## SUPPLEMENT — VIDEOS

**Videos**

**Video 1. Nrp-2 localizes on the plasma membrane of primary cortical neurons in numerous discrete puncta.**

Video 1. pHluorin-Nrp-2 WT_cortical neuron.avi

**Trafficking dynamics of pHluorin-tagged Nrp-2 in primary cortical neurons visualized with FRAP analysis.**

Video 2. FRAP_pHluorin-Nrp-2 WT.avi

Video 3. FRAP_pHluorin-Nrp-2 TCS.avi

